# Multichromatic Dynamic Control of Multi-Membered Microbial Consortia Compositions for Chemical Production

**DOI:** 10.64898/2026.09.28.754827

**Authors:** Jaewan Jang, Emmanouil Alexis, Sebastián Espinel-Ríos, Endri Yzellari, Tiara N. Safaei, José L. Avalos

## Abstract

Engineered microbial consortia offer a promising strategy for chemical production by distributing specialized functions among microbial strains, reducing metabolic burden, facilitating modular pathway optimization, reducing toxicity, and increasing strain stability. However, differences in growth rates can destabilize population composition, compromising productivity and limiting their applicability. Here, we developed a multichromatic optogenetic Toxin-Antitoxin (optogeneticTA) platform for dynamic control of *Escherichia coli* consortia of up to four members using blue, red, and near-infrared light and darkness. By varying light intensities or pulses, we precisely program and dynamically modulate the population composition of two-, three-, and four-membered consortia. We further developed a modular mathematical framework that captures and predicts population dynamics of these optogenetically controlled co-cultures. Applying dynamic control to a two-membered engineered consortium increased phenol production by ∼69% relative to unregulated consortia. These results establish a programmable platform for stabilizing and dynamically optimizing microbial consortia, with potential applications across microbial biomanufacturing.

## Main

Metabolic engineering has enabled the production of a wide variety of chemicals, such as pharmaceuticals, agrochemicals, specialty and commodity chemicals, and biofuels^1^ by rewiring microbial metabolisms. Pathway engineering frequently relies on expressing heterologous genes and deleting endogenous host genes to redirect metabolic flux to a desired product, which often leads to metabolic burden, reduced growth, capped productivity, and ultimately strain instability. Distributing the genetic interventions required to engineer and optimize a metabolic pathway across multiple strains instead of in a single strain has been shown to effectively reduce these unintended side effects.^2–4^ Distributing different modules of a biosynthetic pathway and the required gene deletions across different members of microbial consortia not only lowers the burden in each strain, but also allows optimization of each module independently in each individual strain, as opposed to needing to make compromises that may benefit one module but negatively impact another. Distributed metabolic modules can also help avoid the accumulation of toxic intermediates, and circumvent endogenous regulatory pathways^5–8^. Additionally, microbial consortia can harness the unique capabilities of different microbial species, integrating their specialized functions in a single bioprocess^5,9–11^. Such synthetic microbial co-cultures have been used to produce commodity chemicals such as phenolic derivatives^12–14^ or n-butanol^15^, as well as organic acids^9,16,17^, terpenoid-based^5,18,19^, and flavonoid-based natural products^8,20^, in several cases achieving titers that single-strain-monocultures could not reach^13,14,17,19^.

However, despite many remarkable successes at laboratory scale^21^, engineered co-cultures have not yet been implemented in commercial bioprocesses, with a key challenge being maintaining population stability. If differences in growth rates between consortia members are unaddressed, the faster growing strains can easily displace the slower ones, overtaking the population and turning co-cultures into monocultures of the faster growing strains. Several strategies have been explored to mitigate this challenge. Starting inoculums with a higher proportion of the slower strain or delaying inoculation of the faster one are easy solutions to implement that can reduce the advantage of the faster growing strain^22–24^, but stochastic variations resulting from solely relying on inoculums are difficult to adjust mid-cultivation, which could lead to significant batch-to-batch variations. Tailoring media composition or assigning different carbon sources to individual strains can reduce competition between consortia members^5,9,25^, but careful control of media composition can be difficult to implement in commercial processes. Rational engineering of microbial cooperation via cross-feeding, cross-talk, or co-dependence are elegant solutions to prevent the loss of any consortia member^5,12,26–28^, but the range of accessible consortia compositions by this approach is bound by the emergent dynamics of engineered symbioses, and member proportions can be difficult to control. While these approaches have significantly advanced microbial consortia design for biomanufacturing by improving co-culture stability, even the most sophisticated strategies are limited in their ability to finely, reversibly, and dynamically tune population composition for optimal metabolic coordination, which will likely be necessary for scaling up consortia to commercial processes.

Light regulation through optogenetics offers an attractive strategy to achieve this level of dynamic control over microbial consortia. Optogenetics uses photoresponsive proteins to control biological processes, exploiting many unique advantages of using light as an inducing agent. Light is infinitely tunable, instantly reversible, and easy to manipulate dynamically. It is also inexpensive, orthogonal to biological context, and easily interfaced with computers for closed-loop control and automation^29–31^. Optogenetic systems have been used to control two-membered consortia of *E. coli* using either blue/dark or green/red light-responsive systems to balance the expression of bacterial toxin/antitoxin systems^32,33^, or antibiotic resistance^34,35^. They have also been used to control mixed populations of *S. cerevisiae* by regulating cell differentiation^36^. However, with one exception using only blue light to control *E. coli*-*S. cerevisiae* co-cultures^33^, these light-controlled consortia have not been applied for chemical production.

Here, we describe a multichromatic optogenetic framework for dynamically controlling co-cultures of up to four *E. coli* strains using variable doses of blue, red, and near-infrared (NIR) light, as well as darkness. Building on an optimized blue light-responsive Opto-Toxin/Antitoxin (OptoTA) system that employs pDawn to control MazE/MazF (BlueTA), we developed a new set of optogeneticTA circuits responsive to red light, NIR light, and darkness (RedTA, NIRTA, and DarkTA). Together, these circuits enable orthogonal control of the growth rates of four *E. coli* strains and thus dynamic regulation of population composition across all possible combinations of two-, three-, and four-membered consortia. We further developed a modular modeling framework that quantitatively captures the population dynamics of all four optogenetic strains in both monoculture and multi-strain cultures. We then demonstrate the utility of this approach for bioproduction, showing that light-enabled dynamic control of a two-membered consortium increases phenol production by ∼69% relative to an unregulated consortium. Overall, this study establishes a multichromatic optogenetic platform for dynamic control of *E. coli* consortia comprising up to four members, together with a predictive modeling framework that captures their regulatory behavior. Beyond reproducibly stabilizing consortia, this platform enables population compositions to be precisely programmed and dynamically optimized throughout co-cultivation, providing a framework that could be applied to enhance the production of a broad range of value-added chemicals using microbial co-cultures.

## Results

### Development and characterization of multichromatic optogenetic control over microbial growth

We built multichromatic growth control of *E.* coli from the bacterial MazE/MazF toxin/antitoxin (TA) system^33^ and the light-responsive pDawn/pDusk circuit^37^, which were previously combined as OptoTA, in which the expression of MazE/MazF is photoregulated to grow under blue light while darkness represses it^33^. In darkness, the photoresponsive histidine kinase, YF1, phosphorylates FixJ, inducing *cI* and MazF toxin expression, arresting cell growth (Fig. 1a). Under blue light, YF1 dephosphorylates FixJ, stops the induction of *cI* and MazF expression, and relieves the *cI*-mediated repression of promoter R (pR), thereby driving the expression of MazE antitoxin that inhibits MazF, allowing near wild-type growth (Fig. 1c). Pulsed blue light tunes growth to intermediate levels while fully repressing growth in darkness. To reflect the multichromatic aspect of this study, we renamed this system to BlueTA.

**Figure 1.**
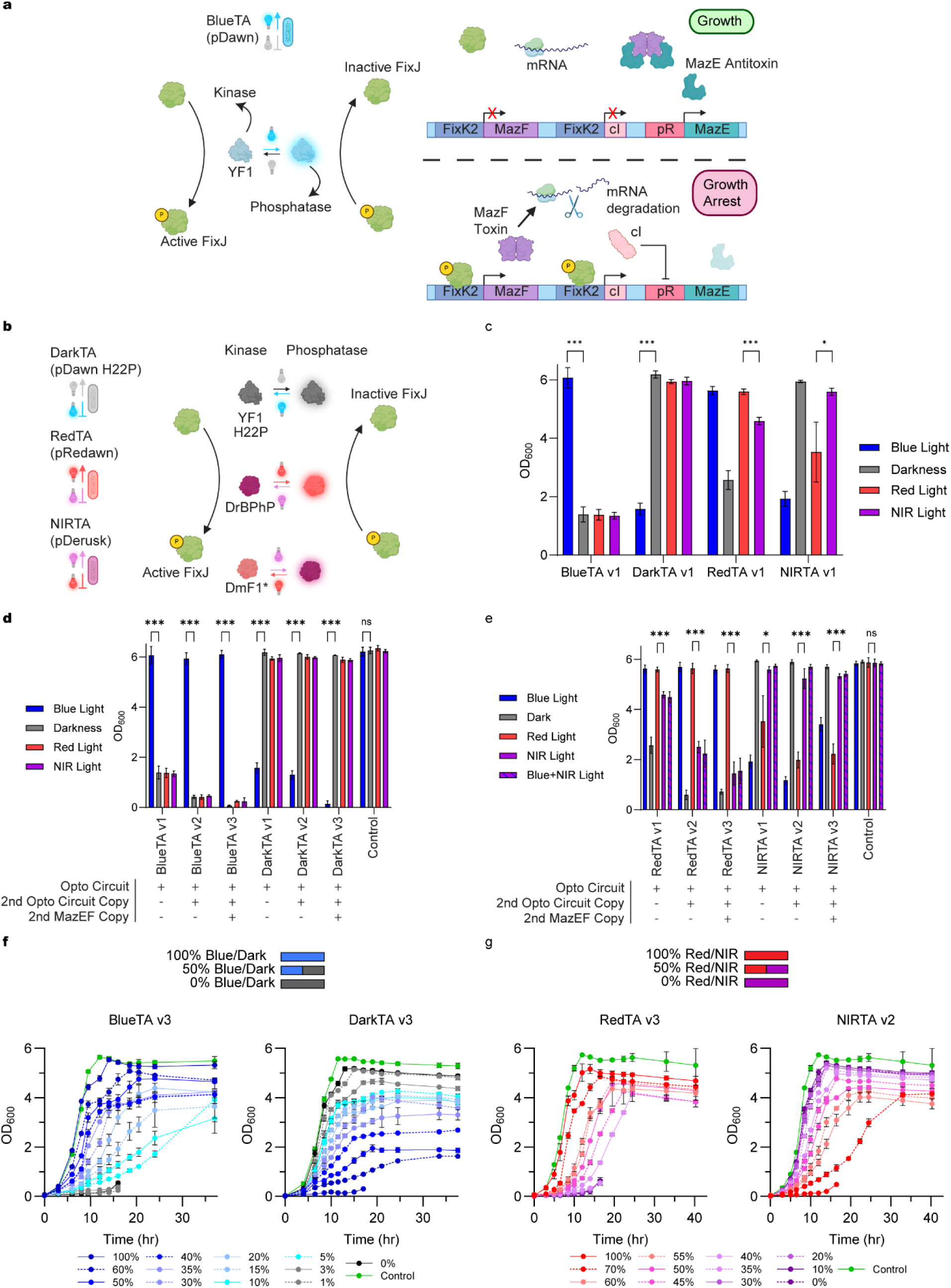
Engineering multichromatic control of optogenetic toxin-antitoxin systems in *E. coli*. **a** Schematic of the BlueTA architecture. A photoresponsive histidine kinase (YF1) drives FixJ phosphorylation or dephosphorylation to activate or repress the MazE-MazF toxin-antitoxin module. Under blue light conditions, YF1 acts as a phosphatase of FixJ, preventing expression of cI repressor and thus enabling expression of MazE antitoxin and cell growth. In darkness, YF1 acts as a kinase that phosphorylates FixJ, which in turn induces MazF toxin activity, triggering mRNA degradation and halting growth. **b** Modular replacement of photoresponsive histidine kinases to afford three additional optogenetically controlled *E. coli* strains. Replacing YF1 with an inverted mutant, YF1^H22P^, or bacteriophytochromes, DrBPhP (pRedawn) or DmF1 (pDmDERusk + *cI*), while leaving the rest of the modules unchanged gives three additional strains that grow either in darkness, red light, or NIR light. **c** Cell growth (OD_600_) of strains BlueTA v1, DarkTA v1, RedTA v1, and NIRTA v1 in red, NIR, blue light, or darkness after 24 hours at 37 °C. **d,e** Cell growth (OD_600_) of (**d**) BlueTA/DarkTA or (**e**) RedTA/NIRTA v1-3 circuits under red, NIR, blue light, and darkness. For RedTA/NIRTA, an additional condition where NIR light co-irradiated with blue light was used to improve orthogonality. Measurement was done after 24 hours at 37 °C. **f,g** Growth tuning of each optogenetic strain under different light-pulse regimes. Growth measurements (OD_600_) of BlueTA v3 (f, left), DarkTA v3 (f, right), RedTA v3 (g, left), and NIRTA v2 (g, right) under 0 to 100% blue light/darkness (**f**) or red/NIR light (**g**) pulses at a forcing period of t_FP_=2000 s. For example, 40% of red/NIR pulse at t_FP_=2000 seconds refers to red light irradiated for 800 seconds (with NIR light off), followed by 1200 seconds of NIR light irradiation (with red light off). Each strain demonstrates robust and predictable growth control using multichromatic light input. For all experiments, blue, red, and NIR light were pulsed or irradiated with intensities of 0.32, 0.17, and 0.28 mW/cm^2^, respectively. Statistics are derived from t-test (***P<0.001, **P<0.002, *P<0.033, ns: not significant). All data are shown as mean values with error bars representing standard deviation from n=3 biological replicates for (c)-(e) and n=4 for (f) and (g).

To develop strains responsive to multichromatic light, we replaced YF1 in the pDawn/pDusk architecture with other FixJ-acting photoresponsive kinases, and tested their ability to multichromatically express GFP (Fig. 1b and Supplementary Fig. 1). Coupling to the *mazEF* OptoTA system created three additional, multichromatic TA systems: DarkTA (grows in darkness, repressed in blue light), RedTA (grows in red light, repressed in NIR light), and NIRTA (grows in NIR light, repressed in red light) (Fig. 1a,b). Both BlueTA and DarkTA exhibited approximately 4-fold greater growth under permissive conditions (blue light and darkness, respectively) and were unresponsive to red and NIR light, demonstrating orthogonality (Fig. 1c). For RedTA, though red light supported the highest growth, NIR illumination still permitted growth at approximately 80% level of red light. By contrast, NIRTA showed approximately 1.6-fold greater growth in NIR light than in red light. Notably, darkness was better than NIR illumination at either suppressing or promoting growth of RedTA or NIRTA, respectively, whereas blue light illumination had similar effects as red light in these systems (Fig. 1c).

We hypothesized that the unwanted growth of optogeneticTA strains under expectedly repressive conditions was due to a combination of circuit leakiness or bacterial escape (“cheating”). To mitigate these effects, we explored adding tightness and redundancy to these circuits since adding regulatory redundancies can be effective at reducing cheating behavior in *E. coli*^38,39^. Therefore, the original strains (v1) transformed with a plasmid containing the original photosensory components of the circuit (FixJ, histidine kinases, and HO when needed) and *mazEF*, were compared to strains transformed with an additional plasmid containing redundant copies of either the photosensory components alone (v2) or both photosensory components and *mazEF* (v3) (Supplementary Fig. 2) while original strains (v1) were transformed with an additional empty plasmid. Indeed, adding a second copy of the photosensory component plus *mazEF* (v3) improved the circuit performance to 78-fold, 43-fold, 3.9-fold, and 2.4-fold for BlueTA, DarkTA, RedTA, and NIRTA, respectively (Fig. 1d,e). For BlueTA, DarkTA, and RedTA, we proceeded to use the v3 version, while for NIRTA, we chose a better-performing v2 one (2.6-fold).

We tested the tunability of the growth rates of these strains by varying light pulses using “forcing period (t_FP_)”, defined as the time interval at which light is pulsed for a certain percentage of time (Fig. 1f-g). For example, 20% blue/dark pulse at t_FP_=1000 seconds indicates 200 seconds of blue light, followed by 800 seconds of darkness while 20% red/NIR pulse at the same t_FP_ indicate 200 seconds of red light, followed by 800 seconds of NIR light. With various blue/dark pulses at t_FP_=2000 sec, BlueTA and DarkTA exhibited tunable growth response (Fig. 1f left and right, respectively) while RedTA and NIRTA responded correspondingly to red/NIR pulses at the same t_FP_ (Fig. 1g left and right, respectively). Notably, after 20 hours, all four optogeneticTA systems exhibited some growth under fully repressive conditions, indicating the cheating behavior even in these improved strain versions (Supplementary Fig. 3&4). It is well established that bacteria often escape regulatory control through silencing mutations or deletion of regulatory genes (in this case, the optogeneticTA circuit), especially in the most restrictive growth conditions, despite having redundant copies of the circuit^40–42^. However, we hypothesized that under less repressive conditions and in the context of microbial consortia, this cheating behaviour would be attenuated (see below).

### Two-membered co-culture population control

To study the behavior of optogenetically controlled consortia, we first established BlueTA/DarkTA co-cultures inoculated at 1:1 initial ratios (total OD=0.1), and incubated in variable blue light pulses to assess their light sensitivity (Fig. 2a). To quantify population compositions, we labelled the BlueTA and DarkTA strains with constitutive expression of mTagBFP2 and sfGFP, respectively, to measure their compositions after 48 hours (Fig. 2b). To decouple pulse timing from intensity, we fixed blue light intensity at a saturating level (∼0.32 mW/cm^2^) and varied pulsing sequence and forcing period (t_FP_). At short t_FP_=200 s, we observed a sharp transition in population composition from DarkTA to BlueTA upon increasing light pulse percentage, reaching a “crossover point” (the light dose at which both strains are present at 50% of the population) at only ∼3.7% blue/dark pulse (Fig. 2b). The high sensitivity indicated that at short t_FP_, a minimal blue light dose was sufficient to switch between BlueTA- and DarkTA-dominated states. The intrinsic thermal relaxation (or dark reversion) time of photoresponsive kinases from the light-activated to dark-adapted states, as well as the half-lives of the mRNA and proteins they induce, are expected to be influenced by the forcing periods and on cell growth. Thus, we hypothesized that longer forcing periods would allow more time for the activated forms of YF1 and YF1^H22P^ to relax as well as induced mRNA and protein levels to more extensively decrease between pulses, thereby shifting the crossover point to higher light doses and smoothening the transition between populations (Supplementary Fig. 6). Consistent with this prediction, extending t_FP_ to 2000 s increased the crossover point to 30%. Further increasing t_FP_ to 4000 and 8000 seconds broadened the dynamic range of compositional control – the range we define as light condition ranges that allow population composition change from 5 to 95% of either strain – and increased the pulsing percentage needed to reach the crossover point (to ∼41% and ∼46%, respectively).

**Figure 2.**
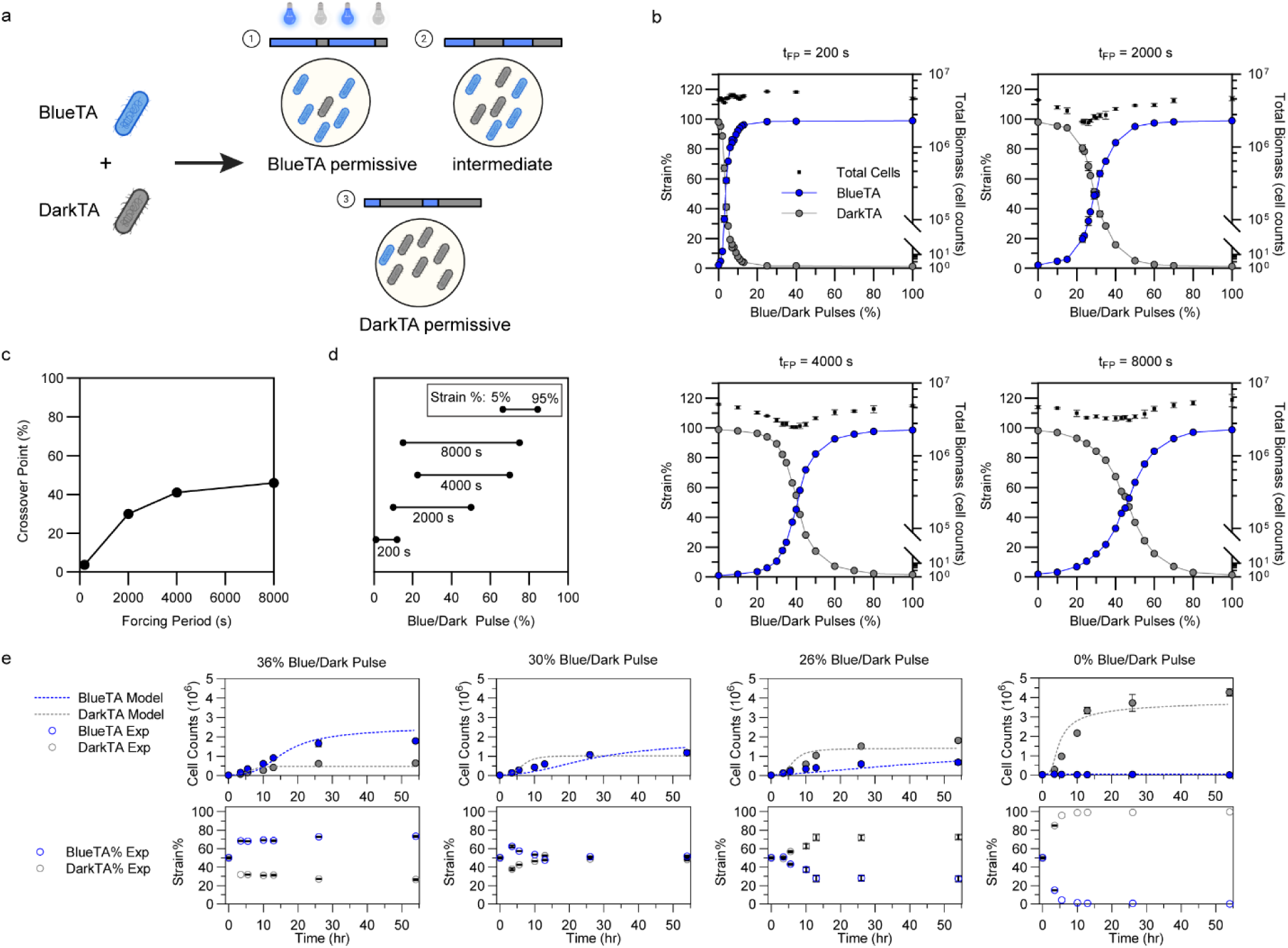
Optogenetic control of two-membered *E. coli* consortia using blue/dark pulses. **a** Schematic illustration of optogenetic population control over a BlueTA and DarkTA consortium. Independent modulation of blue/dark pulse ratios (at 0.32 mW/cm^2^ of blue light) steers the relative abundance of each strain in the co-culture. **b** Final BlueTA and DarkTA strain distributions (in %) across a range of blue/dark pulses under different forcing periods (t_FP_=200, 2000, 4000, 8000 s) after 48 hours at 37 °C. Increasing blue light dose (relative to dark) monotonically increases BlueTA abundance while proportionally suppressing DarkTA. **c** Crossover points, the percentage of blue/dark pulse where the two strains are equally populated (50% strain ratio of each strain), are plotted as a function of forcing period (t_FP_). **d** Dynamic range of light conditions in which the composition of a blue light-controlled consortium is tunable across 5 to 95% population distributions, plotted for different forcing periods (200, 2000, 4000, and 8000 seconds). **e** Time course of BlueTA/DarkTA co-culture growth distributions at selected pulse programs (36%, 30%, 26%, and 0%; from left to right) and t_FP_=2000 s. n% of blue/dark pulse indicates n% of blue light irradiation, followed by (100-n)% of darkness at a certain t_FP_. Top panels show total cell counts by strain, while bottom panels show percent distributions of each strain. Additional pulses tested are in Supplementary Fig. 7. All data are shown as mean values with error bars representing standard deviation from n=3 biological replicates.

We next assessed whether we could control BlueTA/DarkTA population ratios by delivering blue light in the form of intensity as opposed to pulsing (duty cycles). In complete darkness, DarkTA overwhelmingly dominates the culture (>98% of the population), whereas BlueTA reaches >98% of the population at remarkably low blue light levels, indicating that even the minimum intensity provided by the OptoWells (∼0.06 mW/cm^2^) was enough to fully induce the light-dependent growth response of this strain (Supplementary Fig. 5a). To access intensities lower than the minimal emitted by the OptoWells, we used neutral-density filters to attenuate the blue light intensity to ∼5.1% of the emitted value (see Methods). With this adjustment, we were able to tune the composition of co-cultures using intensities ranging from 0 – 36 µW/cm^2^, reaching the crossover point at ∼15 µW/cm^2^ (Supplementary Fig. 5b). Together, these results demonstrate that varying pulsing periods or percentage enables precise control of co-culture composition via the BlueTA/DarkTA co-culture system.

We also combined RedTA/NIRTA strains to form consortia where population ratios were controlled bidirectionally. When using pulsed red/NIR illumination, the crossover point was less sensitive to forcing period than in BlueTA/DarkTA consortia (Fig. 3b,c), at approximately 30.5% red/NIR pulse (t_FP_=200 s), but then stabilizing at ∼49% with t_FP_ ranging from 2000-8000 s, whereas the BlueTA/DarkTA showed a continuous increase in crossover point value with increasing t_FP_ (Fig. 2c). In addition, t_FP_ in the RedTA/NIRTA consortium did not meaningfully affect the dynamic range of compositional control which is stable at ∼29-43% across forcing periods ranging from t_FP_=2,000 to 8,000 s (Fig. 3b,d). In contrast, the dynamic range of compositional control of the BlueTA/DarkTA consortia continuously increased from 11% to 60% with forcing periods ranging from 200 to 8,000 s (Fig. 2d). These observations could reflect the differences in the time it takes to inactivate the light-induced histidine kinase activity at different t_FP_ times (Supplementary Fig. 6). Together, these results show predictable ratio control across a wide range of red/NIR pulsing regimes.

**Figure 3.**
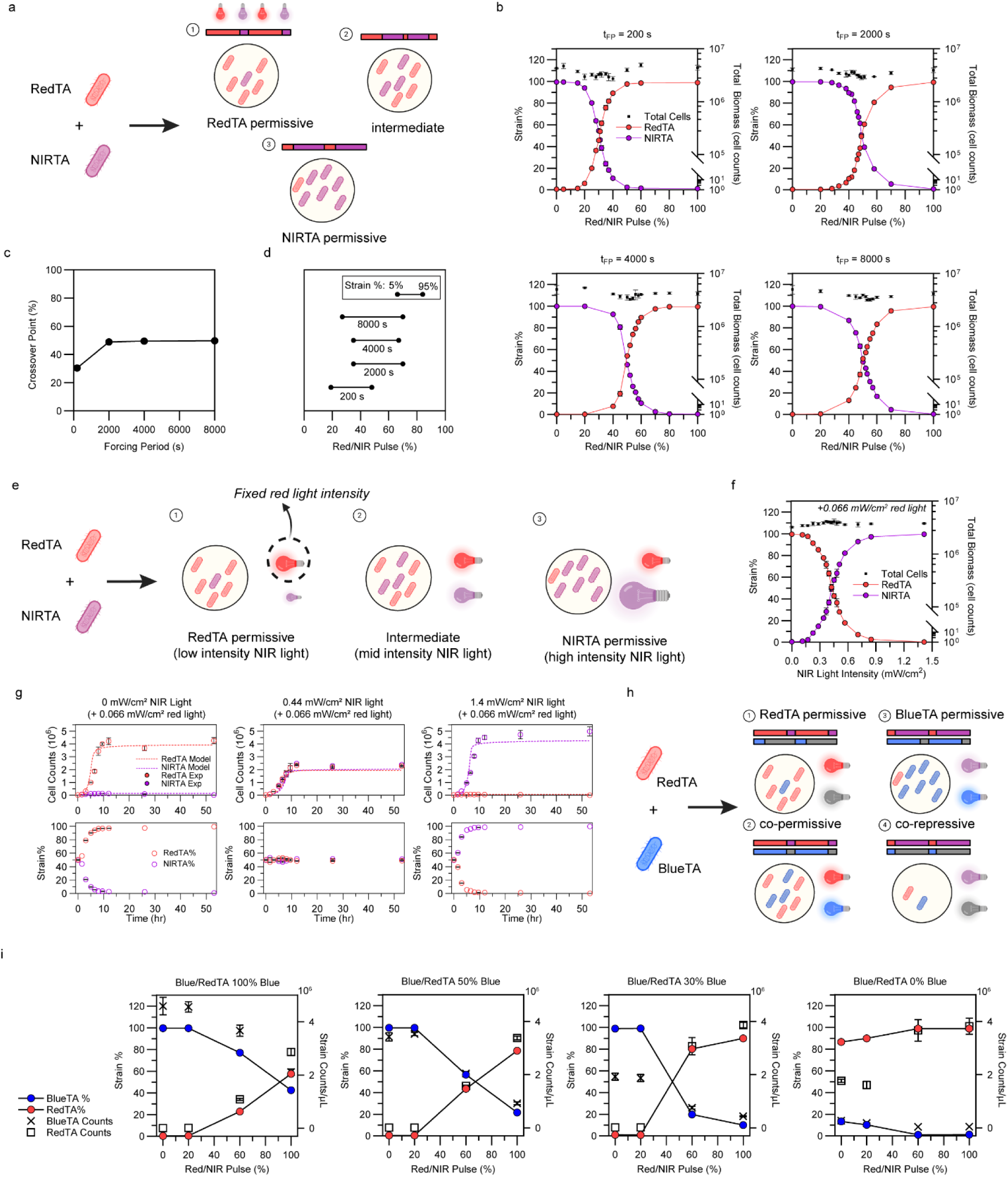
Optogenetic control of two-membered *E. coli* consortia using additional wavelengths. **a** Schematic illustration of optogenetic population control over the RedTA and NIRTA consortium. Independent modulation of red/NIR pulsing percentage steers the relative abundance of each strain in co-culture. **b** Final RedTA and NIRTA strain distributions (in %) across a range of red/NIR pulses under different forcing periods (t_FP_=200, 2000, 4000, 8000 s) after 48 hours at 37 °C. Increasing red light dose (relative to NIR) increases RedTA proportion while proportionally suppressing NIRTA growth, and vice versa. n% of Red/NIR pulse indicates n% of red light irradiation, followed by (100-n)% of NIR light irradiation at a certain t_FP_. **c** Crossover points, the percentage of red/NIR pulse where the two strains are equally populated at 50% strain ratio, are plotted as a function of forcing period (t_FP_). **d** Dynamic range of light conditions in which the composition of a red and NIR light-controlled consortium is tunable across 5 to 95% population distributions, plotted for different forcing periods (200, 2000, 4000, and 8000 seconds). **e** Schematic illustration of optogenetic population control over the RedTA/NIRTA consortium by light intensity modulation. Under fixed red light intensity, varying NIR light intensity modulates the relative abundance of the two strains. **f** Effect of modulating NIR light intensity (while keeping a fixed red light background, 0.066 mW/cm^2^) on RedTA/NIRTA population composition after 48 hours at 37 °C. Increasing NIR intensity suppresses RedTA and enhances NIRTA growth affording predictable consortia population compositions. **g** Time course of RedTA/NIRTA co-culture compositions grown under constant red light (0.066 mW/cm^2^) and additional exposure to modulated intensities of counteracting NIR light (at selected intensities of 0, 0.44, and 1.4 mW/cm^2^). Each strain converges to the expected population composition based on counteracting NIR light intensity used. The RedTA/NIRTA consortia model, adjusted to account for simultaneous light-intensity illumination, effectively captured the overall growth dynamics. Top panels show total cell counts by strain, while bottom panels show percent distributions of each strain. Additional intensities tested are in Supplementary Fig. 9. **h** Schematic illustration of a multi-dimensional consortia population control strategy combining BlueTA and RedTA strains. By independently modulating blue/dark and red/NIR pulses, growth of BlueTA and RedTA can be simultaneously and orthogonally tuned. **i** Final RedTA and BlueTA strain % across a range of Red/NIR pulses at four different blue/dark pulses (100, 50, 30, 0%) after 24 hours at 37 °C with the co-culture inoculated at 1-to-1 ratio. While increasing blue light dose promotes BlueTA growth, increasing red/NIR pulse suppresses it due to simultaneous growth of the RedTA strain. Under simultaneous growth-repressed conditions, a small fraction of RedTA growth is observed due to the emergence of cheater cells. For pulsing-based experiments, blue, red, and NIR light were pulsed with intensities of 0.32, 0.17, and 0.28 mW/cm^2^, respectively. All data are shown as mean values with error bars representing standard deviation from n=3 biological replicates.

We next assessed how the RedTA/NIRTA consortium responds to varying light intensities as opposed to pulsing. Both RedTA and NIRTA strains were highly sensitive to light intensities as low as 32.6 µW/cm^2^ red light and 28.4 µW/cm^2^ NIR light, respectively, giving >98% of the corresponding strains after 48 hours, limiting the dynamic range of tunability (Supplementary Fig. 5c-d). To mitigate this high sensitivity and extend the dynamic range to control consortia composition, we simultaneously illuminated with red and NIR light by varying intensities of each color (Fig. 3e); thus, the two wavelengths act as opposing actuations on the photoresponsive histidine kinases – enabling more precise tuning of the co-culture composition. When red light intensities were fixed at 0.066 mW/cm^2^, progressively increasing NIR light intensity from 0 to 1.4 mW/cm^2^ shifted the population towards the NIRTA strain (Fig. 3f). Time course experiments showed that with NIR light intensity tuning, the co-cultures rapidly approached final ratios at 7.5 hours after inoculation (Fig. 3g and Supplementary Fig. 9). At half red light intensity (0.033 mW/cm^2^), half the intensity of NIR light was needed to reach similar population ratios (Supplementary Fig. 10a), and the final strain ratio obtained at a given red/NIR light intensity ratio is approximately the same as the overall light intensity increases (Supplementary Fig. 10b). Thus, in contrast to the BlueTA/DarkTA system, which is controlled by monochromatic intensity (Supplementary Fig. 5a&b), the RedTA/NIRTA system enables the balancing of red/NIR intensity ratios.

Illumination regimes also influence total biomass accumulation across the multicolor co-cultures. When controlling consortia with light pulses, with the exception of BlueTA/DarkTA at t_FP_=200 s, both types of consortia show slightly reduced final total biomass accumulation when exposed to intermediate levels of light pulses (with minima near the crossover points) (Fig. 2b, 3b). Under simultaneous intensity-modulated red/NIR illumination, intensity ratios giving intermediate population ratios yielded relatively similar final biomass across all growth conditions of both RedTA and NIRTA (Fig. 3f and Supplementary Fig. 10a). Thus, intensity-modulated control not only enables precise compositional tuning but also preserves overall biomass yield across different light stimuli – an advantage over pulse-modulated control, especially for applications where total community biomass is critical like in chemical production.

In the BlueTA/DarkTA or RedTA/NIRTA consortium, light inputs always stimulate both members of the consortia in opposite directions (e.g., in red light the RedTA strain grows while the NIRTA is inhibited, and vice versa in NIR light), which could slow overall growth of the communities. We paired RedTA with either BlueTA or DarkTA to make the growth response orthogonal to each strain, enabling co-permissive growth under combined blue, red, and NIR inputs (Fig. 3h and Supplementary Fig. 11a). In RedTA/BlueTA co-cultures under fully permissive illumination (red plus blue light), both members stabilized at similar levels (60/40%, respectively) with overall biomass comparable across different pulses, consistent with maximizing growth of both strains simultaneously, albeit each one having inherent differences in their maximal growth rates (Fig. 3h,i). Reduction of red/NIR light pulse shifted the culture towards BlueTA growth, indicating that RedTA abundance remained tunable by red and NIR input even when BlueTA was fully activated. Conversely, reduced blue-light illumination constrained BlueTA growth and lower red/NIR pulse (lower and higher dosage of red and NIR light, respectively) was required to approach crossover points, consistent with its strict blue-light dependence. Under co-repressive NIR-only illumination, both strains are repressed, though some cheating behavior can be observed as in the monoculture experiments. The RedTA/DarkTA consortium showed analogous wavelength-dependent behavior (Supplementary Fig. 11b-e). Under red-only illumination, both strains grew permissively, whereas decreasing red light input (increasing NIR input) shifted the culture toward DarkTA dominant growth. Blue-light dosing restricted DarkTA growth while RedTA remained responsive to red/NIR light modulation.

Similar wavelength-dependent responses were observed when NIRTA was paired with BlueTA or DarkTA (Supplementary Fig. 12), with each consortium showing predictable and reversible population shifts in response to blue, red, and NIR inputs, demonstrating multicolor capabilities of the optogeneticTA system in two-membered consortia of all possible combinations (Fig. 2&3, and Supplementary Fig. 11&12).

### Three- and four-membered co-culture control

To demonstrate optogenetic control over communities with more than two-members, we first combined the BlueTA strain with RedTA and NIRTA in three-membered co-cultures (Fig. 4a). Under fixed red/NIR light pulses at 48%, 50%, and 52%, the RedTA/NIRTA ratio stabilized at ∼55/45%, 50/50%, and 40/60%, respectively (Fig. 4a), while increasing blue/dark pulses tuned the BlueTA fraction from 0-60%, approximately. Importantly, the relative RedTA/NIRTA ratio was unaffected by blue light modulation due to the saturating red and/or NIR light intensities that offset the influence from blue light. Time-course measurements confirmed that the population ratios rapidly converged to the desired composition under fixed 50% red/NIR and blue light pulses (Fig. 4b and Supplementary Fig. 13). The same predictable behavior and capabilities were observed when combining the DarkTA strain with RedTA and NIRTA strains (Supplementary Fig. 14, 15), as well as when combining RedTA or NIRTA with BlueTA and DarkTA (Supplementary Fig. 16).

**Figure 4.**
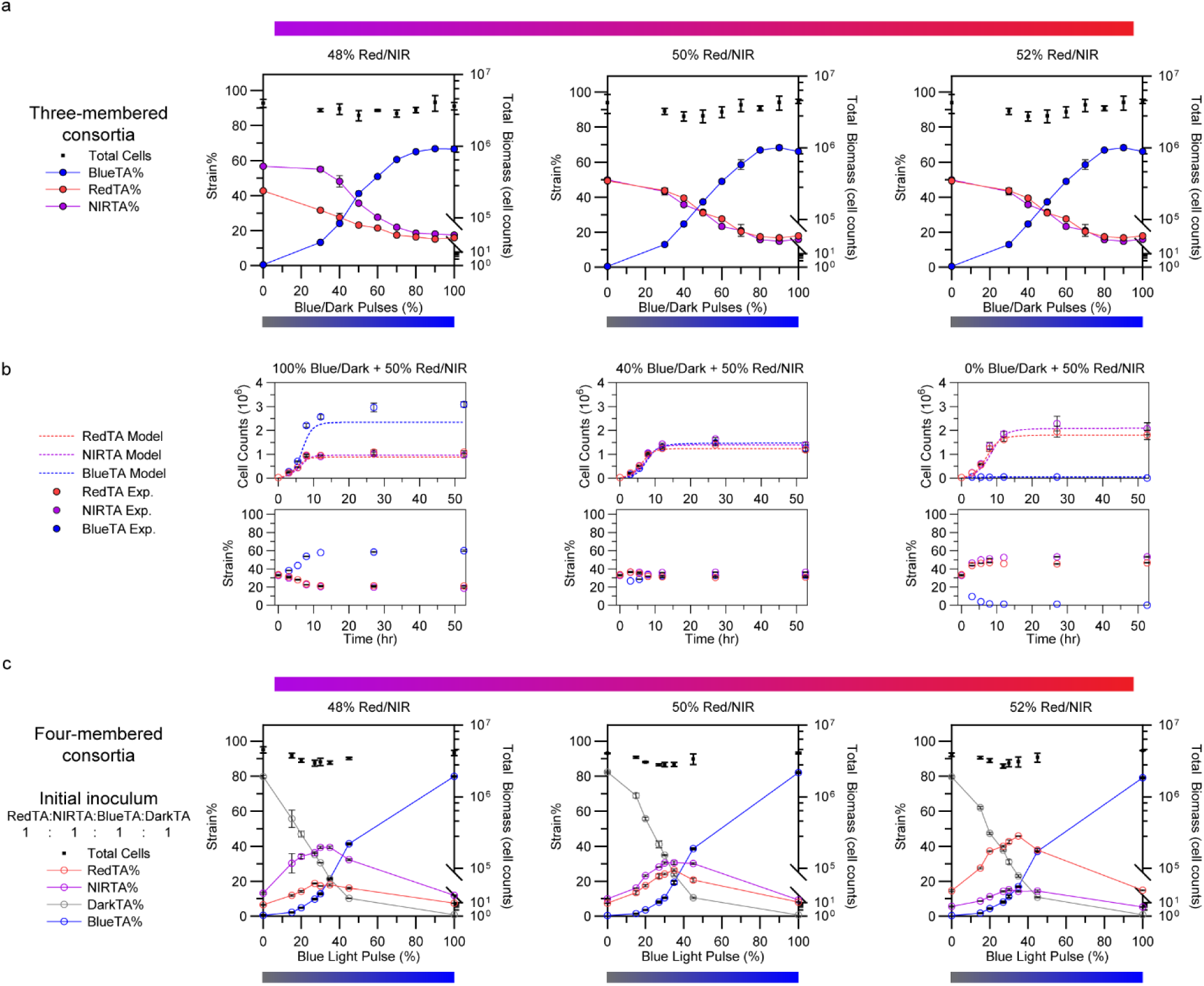
Orthogonal multichromatic optogenetic control of three- and four-membered consortia. **a** Final population distributions of BlueTA/RedTA/NIRTA co-cultures incubated at 37 °C for 48 hours at three different red/NIR light pulses (48, 50, and 52%; left to right) and varying blue/dark pulses (t_FP_=2000 seconds) after 48 hours at 37 °C. BlueTA composition responds solely to blue/dark pulses while RedTA/NIRTA population ratios are set by the red/NIR pulses. n% of Red/NIR pulse indicates the n% of red light irradiation, followed by (100-n)% of NIR light irradiation at a certain t_FP_. **b** Time course of population composition of BlueTA, RedTA, and NIRTA under representative 100, 40, and 0% blue/dark pulses (left to right) while co-irradiated with 50% red/NIR pulse. Top panels show cell counts of each strain as measured from the flow cytometer with model fit indicated in dashed lines. Bottom panels show co-culture compositions by strain percentage obtained from strain counts. All strains converge to specific compositions in response to different multichromatic light input. Additional pulses tested are in Supplementary Fig. 13. **c** Final population distributions of the BlueTA/DarkTA/RedTA/NIRTA four-membered consortium inoculated with 1:1:1:1 ratio, respectively, incubated at 37 °C for 48 hours at fixed red/NIR pulses (48, 50, and 52%; left to right) and varying blue/dark pulses (t_FP_=2000 s). Additional inoculation ratios tested are in Supplementary Fig. 18. In all experiments, blue, red, and NIR light were pulsed at 0.32, 0.17, and 0.28 mW/cm^2^. All data are shown as mean values with error bars representing standard deviation from n=3 biological replicates.

The composition of a three-membered optogenetically controlled consortium can also be fine-tuned using intensity modulation. While red and NIR light intensity modulation reliably adjusts the RedTA/NIRTA ratio, we found that continuous blue light illumination in three-membered systems introduced modest perturbations to this ratio (Supplementary Fig. 17a,b). This effect was expected, as blue-light photons can influence red/NIR-responsive strains. This is consistent with monoculture experiments (Fig. 1d), where blue light stimulated the RedTA strain growth while inhibiting the NIRTA strain, effectively acting as a surrogate for red light. In a three-membered consortium, when blue light was pulsed at 0.13 or 0.32 mW/cm², the extra blue light significantly favored RedTA strain growth over the NIRTA strain. Consistently, this effect is absent under 0% blue illumination and increasingly pronounced at higher blue-light intensity pulses (Supplementary Fig. 17a,b). This response supports a direct, photon-dose-dependent influence of blue light stimuli on RedTA and NIRTA strains.

Because NIR illumination can counteract blue light interference into these red/NIR circuits, we derived an empirical compensation rule to correct for unintended blue-light stimulation of RedTA and NIRTA strains: each 0.13 mW/cm^2^ increase in blue light intensity can be offset by ∼ 0.03 mW/cm^2^ units of NIR (Supplementary Fig. 17c). For example, in a three-membered consortium (RedTA/NIRTA/BlueTA), adding blue light pulses at 0.13 mW/cm^2^ deviated the RedTA/NIRTA in favor of RedTA as more blue light was pulsed (Supplementary Fig. 17d). However, during each blue light pulse, an additional pulse of NIR light at 0.03 mW/cm^2^ intensity helps offset this blue light effect (Supplementary Fig. 17e), keeping the RedTA/NIRTA ratios more consistent across different blue light pulses. Although minor deviations were still observed under extreme regimes (e.g. 100% blue pulses), this can be addressed through future compensation rule optimization or implementation of real-time feedback control.

Finally, we demonstrated a four-membered consortium consisting of all four optogeneticTA strains. To test the response of the four-membered consortium to multichromatic light inputs, we fixed three different red/NIR pulses and varied blue light/dark pulses to modulate the BlueTA/DarkTA ratio (Fig. 4c). As expected, the RedTA/NIRTA maintained a roughly equal ratio at each red/NIR pulsing regime while the BlueTA and DarkTA strain was permissive in blue light and darkness, respectively, across all experiments. We reached a 3-member crossover point at 48% Red/NIR light pulse and ∼35% blue light, and were close to reaching other 3- and 4-member crossover points under other light conditions. Changing the initial inoculation ratios greatly affects the final population ratios, and the conditions in which different multi-member crossover points are achieved (Supplementary Fig. 18a,b), providing a practical strategy for balancing intrinsic growth differences when using more than two consortia members. Together, these results demonstrate an orthogonal and quantitatively programmable control over four-membered microbial co-cultures using multichromatic optogeneticTA regulation.

### Dynamic control of co-culture populations

A key advantage of optogenetics is the ability to dynamically modulate biological systems with orthogonal wavelengths of light. Therefore, we explored how dynamic light inputs could be used to precisely regulate population composition during co-culture growth. To test this capability, we first used two-membered RedTA/NIRTA strain co-cultures, and studied their population dynamics upon shifting light conditions. We started with RedTA/NIRTA co-cultures (inoculated at a 1:1 ratio) illuminated continuously with either 100% red or NIR light. After three hours, we shifted to different proportions of red plus NIR light by varying either light pulses or intensity. When controlling with light pulses and shifting from red to NIR light (Fig. 5a and Supplementary Fig. 19), we found, as expected, that the RedTA strain takes over the population, reaching ∼88% at the initial three hours, and depending on the extent to which light was shifted from red to NIR, the population composition responded more or less rapidly and achieved a larger change in the final consortia composition (Supplementary Fig. 19). For example, when shifting to 40% red, the NIRTA strain population begins to increase 4.5 hours after the light shift, reaching ∼44% of the co-culture after ∼48 hours (Fig. 5a left). In contrast, when shifting completely to NIR light (0% red light), the NIRTA population begins to be favored almost immediately, reaching ∼88% of the population after the same time period (Fig. 5a right).

**Figure 5.**
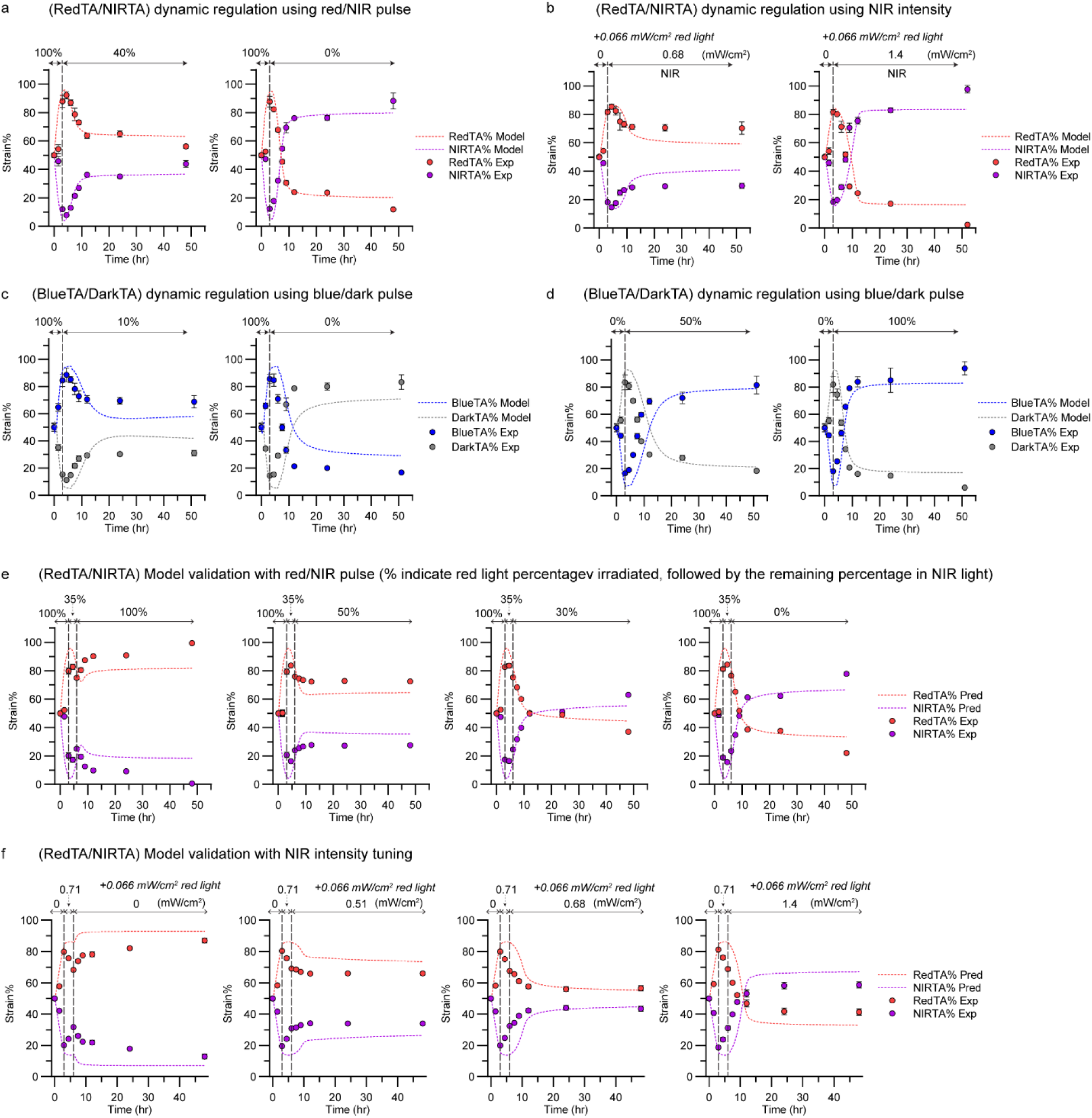
Dynamic control of co-culture populations using multichromatic optogeneticTA systems and model validation (dotted lines). **a,b** Time-courses of RedTA/NIRTA co-cultures in response to dynamic red/NIR pulsing (a) or red/NIR intensity modulation (b). Co-cultures inoculated at 1-to-1 RedTA/NIRTA ratios were first exposed to 100% red light (at 0.17 mW/cm^2^) in (a); or 0 mW/cm^2^ NIR light with 0.066 mW/cm^2^ red light in (b) for 3 h, followed by switching light conditions to the indicated red/NIR light pulsing regimes (a) – 40% red/NIR light (left) at t_FP_=2000 s (800 seconds of red light and 1200 seconds of NIR light of red and NIR light at 0.17 and 0.28 mW/cm^2^, respectively) or 0% red/NIR light at t_FP_=2000 s (0 seconds of red light and 2000 seconds of NIR light at the same intensity) – or the indicated NIR light intensities (b). n% of Red/NIR pulse indicates n% of red light irradiation, followed by (100-n)% of NIR light irradiation at a certain t_FP_. Additional pulses and intensities are in Supplementary Fig. 19 and 20. **c,d** Time-courses of BlueTA/DarkTA co-cultures in response to dynamic blue light inputs. Co-cultures inoculated at 1-to-1 BlueTA/DarkTA ratios were initially exposed to either 100% blue light (at 0.32 mW/cm^2^) (c) or 0% blue light (darkness) (d) for 3 h, followed by switching light conditions to the indicated blue light/darkness pulsing cycles at the same blue light intensity and t_FP_=2000 s. **e,f** Model validation of RedTA/NIRTA co-culture dynamics under different light programs. **e** Cells were exposed to 100% red light (t_FP_=2000 s; 2000 seconds red light only at 0.17 mW/cm^2^) for 3 h, then light conditions were switched to 35% red/NIR light (700 seconds of red light at 0.17 mW/cm^2^ and 1300 seconds of NIR light 0.28 mW/cm^2^; t_FP_=2000 s) for 3 h, followed by a second switch in light conditions to the indicated pulsing schedules (same intensities and forcing periods). **f** Same scheme as in (e) but modulating NIR light intensity instead of pulsing. In all panels, points represent the experimental measurements, and dotted lines indicate either model predictions or model fitting. Dashed vertical lines indicate changes in light pulsing regime or intensity. All data are shown as mean values with error bars representing standard deviation from n=3 biological replicates.

The same trends are observed when dosing the red/NIR light inputs by intensity ratios of the two lights (Fig. 5b and Supplementary Fig. 20). When irradiating co-cultures with only red light at 0.066 mW/cm^2^ for three hours, the RedTA strain reached ∼81% of the composition. Subsequently, co-irradiating NIR at different intensities, without changing red light intensity, rapidly reversed the trend, enriching the NIRTA population and shifting the final RedTA/NIRTA population ratios (Supplementary Fig. 20); for example, to 70% of RedTA strain when co-irradiating with 0.68 mW/cm^2^ NIR (Fig. 5b left) or to 2% RedTA with 1.4 mW/cm^2^ NIR (Fig. 5b right). Similar to the behavior in co-cultures controlled with pulses, the higher the NIR light intensity, the more rapid and dramatic is the population shift.

If co-cultures are first irradiated with NIR light and then, after three hours, exposed to increasing red light stimulation, the same trends but in opposite direction are observed, whether inputs are dosed by light pulses or intensity (Supplementary Fig. 21&22). Similar dynamic controls of BlueTA/DarkTA co-culture populations can be achieved by dosing blue light pulses (Fig. 5c&dand Supplementary Fig. 23). Together, we demonstrate that both pulsed and continuous illumination can provide real-time, dynamic regulation of co-culture composition.

### Modeling

To quantitatively describe and predict the behavior of our light-controllable strains, we developed a modular ordinary differential equation (ODE)-based mathematical framework. For each strain, the model tracks the photoswitching dynamics of the YF1 histidine kinase between its dark- and light-adapted states, the phosphorylation-dephosphorylation cycle of the response regulator FixJ, *cI*-mediated transcriptional regulation, and the toxin-antitoxin (mazF/mazE) interaction that couples gene expression to light-dependent modulation of cell growth. Strain-specific light responses are captured through photoactivation and dark-recovery rate functions that depend on blue, red, or NIR light intensity according to the strain identity (BlueTA, DarkTA, RedTA, or NIRTA). Cell growth is modeled as a nutrient- and mazF-dependent process, enabling the framework to reproduce population-level dynamics under varying illumination conditions (Supplementary Fig. 3&4). The single-strain model is naturally extended to microbial consortia by coupling individual strain sub-models through a shared nutrient pool. For simplicity, the framework does not account for phototoxicity effects or other potential inter-strain interactions such as metabolic cross-feeding or competition beyond nutrient depletion. Upon suitable parameter fitting to experimental time-course data (see also Methods), the model successfully captures the observed macroscopic growth dynamics of both individual strains and multi-strain consortia across a range of illumination conditions (Fig. 2–5 and Supplementary Fig. 7-9, 13, 15, 19-23). Detailed discussion on the model structure, parameter definitions, and fitting procedure can be found in Supplementary Discussion 1.

To test the predictive power of our models, we compared model predictions of various population trajectories for two-membered consortia with experimental measurements. Trajectories with three light irradiation phases were implemented (0-3 h, 3-6h, 6-48h) using variable red/NIR light pulses or intensities to control the RedTA/NIRTA consortium. Despite slight deviations, the model predictions closely followed the overall trends measured experimentally (Fig. 5e,f). Similarly, the model predicted trajectories for the BlueTA/DarkTA consortium demonstrated good agreement with the experimental data under different blue/dark light pulses applied over the three illumination phases (Supplementary Fig. 24). The general applicability of these models to predict multiwavelength light-input trajectories of polychromatically controlled microbial communities unseen during parameter estimation highlights their potential for integration into advanced model-based optimization and control frameworks. Therefore, mathematical modeling was able to capture the overall population dynamics under dynamically varied light pulses.

### Dynamic control of co-culture composition for improving phenol production

Finally, we applied the optogeneticTA systems in combination with metabolic engineering to improve chemical production by microbial consortia through precise multichromatic control over their population composition. As proof of concept, we applied the optogeneticTA control to a microbial consortium engineered for phenol production, which was previously shown to be superior to monocultures^12^. Similar to a previous design^12^, we constitutively expressed a set of six enzymes involved in tyrosine biosynthesis (tyrA^fbr^, aroG^fbr^, aroE, aroL, aroA, aroC) from the proD promoter in an upstream *E. coli* strain (BL21·DE3ΔtyrRΔmazEFΔrecA). Separately, in a downstream strain (MG1655·DE3ΔendAΔrecAΔmazEF), we expressed tyrosine-phenol lyase from *Pasteurella multocida* using the same constitutive promoter (Fig. 6a). We first screened upstream-to-downstream inoculation ratios of the non-optogenetically controlled co-culture (19-to-1, 9-to-1, 3-to-1, 1-to-1, 1-to-3, 1-to-9, 1-to-19) to estimate the ratio giving the highest phenol titer. As expected and previously observed^12^, extremely higher upstream strain inoculum ratios (19-to-1 or 9-to-1) resulted in a larger excess of unconverted tyrosine, whereas a 3-to-1 inoculum ratio gave near-complete conversion of tyrosine to phenol (152 mg/L; Fig. 6b). On the other hand, higher downstream strain inoculation ratios (1-to-3, 1-to-9, and 1-to-19) produced less tyrosine, and thus less phenol.

**Figure 6.**
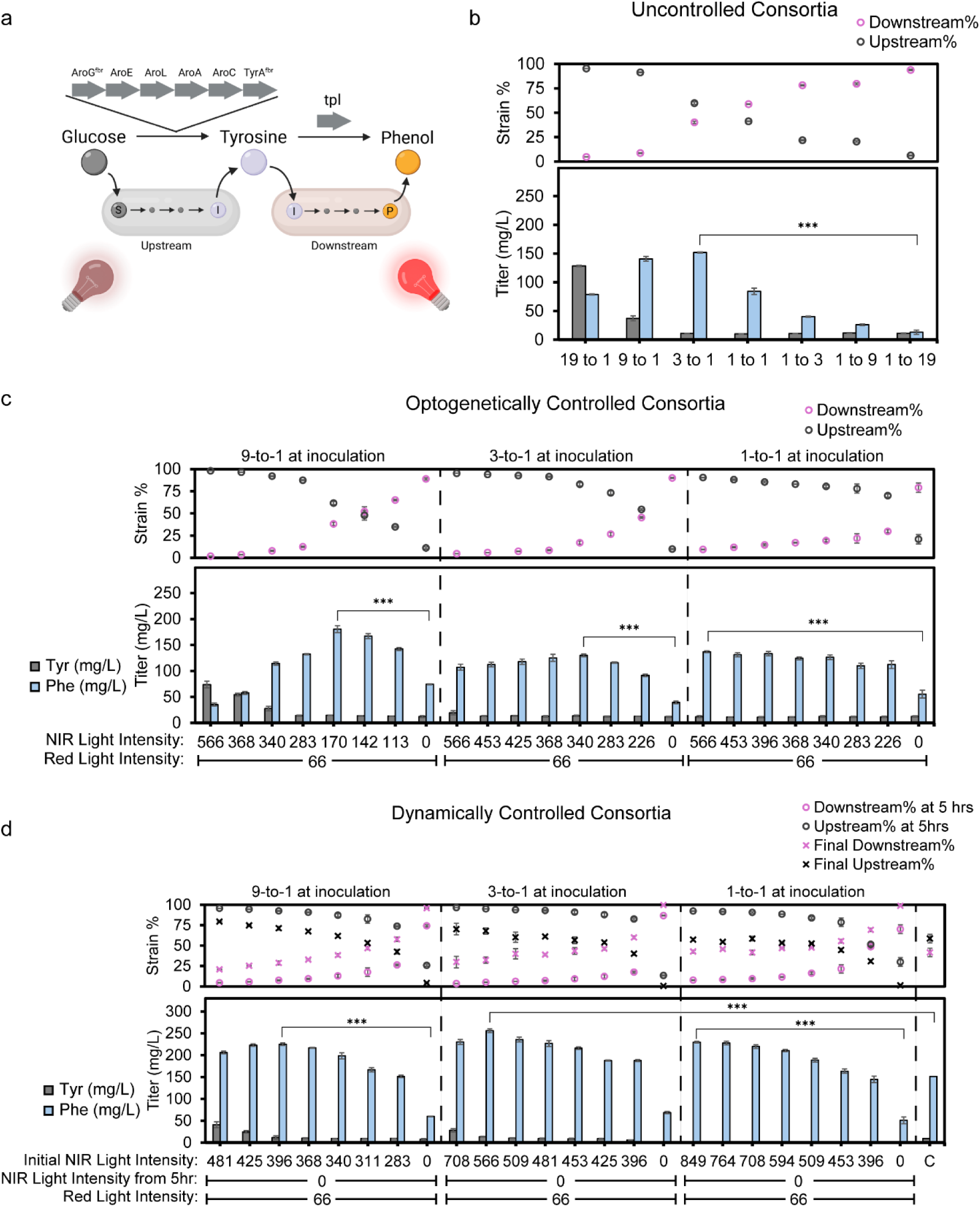
Dynamic control of microbial consortium designed for phenol production using optogeneticTA systems. **a** Schematic of the *E. coli*-*E. coli* consortium in which an upstream strain is engineered to produce and secrete tyrosine, and a downstream strain is engineered to uptake this intermediate and convert it to phenol as the final product. The upstream strain was engineered with the NIRTA optogenetic growth control and six enzymes involved in the shikimate pathway to produce tyrosine, while the downstream strain was engineered with the RedTA optogenetic growth controls and tyrosine-phenol lyase (tpl) to produce phenol from the tyrosine produced by the upstream strain. **b** Optimization of inoculation ratios of upstream:downstream strains in co-cultures lacking optogenetic controls (without NIRTA and RedTA systems) under ambient light. Top panel shows final upstream-to-downstream ratios (after 48 hours), while the bottom panel shows tyrosine and phenol titers (mg/L) of each co-culture. **c** Co-cultures of optogenetically controlled consortia for phenol production starting with different inoculation ratios of upstream/downstream strains controlled with NIRTA and RedTA, respectively. Following inoculation, co-cultures were constantly irradiated with red light at 66 µW/cm^2^ and different NIR light intensities as indicated. Top panels show final upstream-to-downstream strain ratio (after 48 hours), while the bottom panels show the final tyrosine and phenol titers (mg/L). **d** Dynamic control of the population composition of co-cultures for enhanced phenol production using RedTA and NIRTA optogenetic circuits. Starting at different upstream:downstream strain inoculation ratios, co-cultures were irradiated for the initial 5 hours with constant red light (at 66 µW/cm^2^) and different NIR light intensities. Then, the NIR light was switched off while keeping 66 µW/cm^2^ red light constant for the rest of the cultivation. Top panels show both the population ratios of upstream:downstream strains at the 5-hour point (before light switching), and the final population ratios of these strains at the end of cultivation (48 hours). Bottom panels show tyrosine and phenol titers (mg/L). “C” indicates the unregulated consortium (lacking optogenetic control) at the optimal 3-to-1 inoculum ratio under ambient light (b). All phenol and tyrosine titers are indicated in means (mg/L) and the error bars represent standard deviation. Strain percentages are calculated as mean from strain percentages obtained from cell counts on the flow cytometer and the error bars represent standard deviation. Statistics are derived from t-test (***P<0.001, **P<0.002, *P<0.033, ns: not significant).

To encode polychromatic optogenetic controls in this phenol-producing consortium, we introduced the NIRTA system to the tyrosine-producing upstream strain and the RedTA system to the phenol-producing downstream strain (Fig. 6a). Co-cultures were inoculated at upstream:downstream ratios of 9-to-1, 3-to-1, and 1-to-1 while constantly irradiated with red light at a fixed intensity (66 µW/cm^2^) and varying NIR light intensity inputs from 0 to 566 µW/cm^2^, aiming to achieve a similar range of final population ratios across all three inoculums (See *Two-membered Co-culture Population Control*) (Fig. 6c). Cultures inoculated at 9-to-1 and irradiated with high NIR light intensities (566 or 368 µW/cm^2^) left a significant amount of unconverted tyrosine (∼74 or 54 mg/L, respectively), consistent with the unregulated co-culture experiments. At intermediate intensities of NIR light, phenol titer reached the maximum value of 181 mg/L (at 170 µW/cm^2^ NIR) for consortia cultivated at fixed light regimes, leaving only a trace amount of leftover tyrosine, ∼15 mg/L.

A key advantage of optogenetic control over consortia composition is that their populations can be manipulated dynamically by varying light conditions during cultivation, with the potential to further improve chemical production. We reasoned that phenol titer could improve by starting with light conditions that favor the upstream strain growth and then switching mid-cultivation to conditions that favor the downstream strain growth. We tested this hypothesis using three different inoculation ratios, irradiating the co-cultures with variable NIR light intensity (to stimulate the upstream strain), and fixed red light at 66 µW/cm^2^. After 5, 7, or 9 hours, we switched off the NIR light (disfavoring the upstream strain) while keeping red light irradiation unchanged (66 µW/cm^2^) to favor the downstream strain (Supplementary Fig. 25, 26). This resulted in dramatic shifts in population dynamics, from the time of change to the end of cultivation, between the different dynamic and fixed light conditions. In general, the later the light switch occurs, the smaller the change in population ratios relative to static light conditions. For example, with NIR irradiation at 392 µW/cm^2^, the population ratio was kept at roughly 90% of upstream strain, and switching NIR light to 0 µW/cm^2^ at 5, 7, and 9 hours gave upstream strain compositions of 51, 80, and 85%, respectively, while fixed light irradiation mostly kept the ratio unchanged at 90%. In every case, dynamic switching improved the phenol titer compared to fixed illumination, most notably at the 9-to-1 inoculation ratio condition, where most tyrosine was converted to phenol, reaching 212 mg/L, when switching at 5 hours with initial NIR irradiation at 392 µW/cm^2^. The extent to which phenol titer improved depended on the switching time, the initial NIR light intensity, and inoculation ratio. Because switching light conditions at 5 hours was optimal in our set of experiments, we next varied the initial NIR light across a wider range of intensities, while keeping the same time of switching to further improve the titer. Varying this initial irradiation from 283 to 849 µW/cm^2^ resulted again in variable population dynamics (Fig. 6d). At these conditions, starting with a 3-to-1 inoculation ratio, and switching from 566 µW/cm^2^ NIR light to red-light-only at 5 hours gave the maximal phenol titer, 256 mg/L, which is a 69% improvement over the unregulated 3-to-1 co-culture. This titer exceeds the highest reported phenol titer (∼210 mg/L) in batch and using an optimized biosensor-assisted co-culture cultivation strategy^12^.

## Discussion

Microbial consortia offer clear advantages for chemical production, but they have seen little practical deployment due to the inability to stabilize and control their microbial composition. Tuning initial inoculation ratios sets the starting composition but cannot be easily corrected mid-cultivation. This is critical because specific co-culture compositions and their dynamics have a profound impact on titers, yields, productivities, and byproduct formation. While significant progress has been made in improving consortia stability (avoiding co-cultures from drifting towards dominance by the member(s) that grow(s) the fastest), less has been done to achieve tight dynamic control of co-culture composition^12,26,34,43^. Timing the inoculation of different members can afford some dynamic control over consortia composition^26^, but relying on inoculation ratios alone are difficult to tune, irreversible, and prone to stochastic variability between batches, which would be particularly problematic in scaleup applications for chemical production.

Optogenetics is ideally suited for achieving not only consortia stability but also obtaining tight and reversible dynamic control of their strain composition throughout the co-culture. Optogenetic controls of microbial co-cultures have been previously reported for *E. coli*-*E. coli*^32,34^, *E. coli*-yeast^33^, and yeast-yeast consortia^36,44,45^. However, they have important limitations, including controlling only one member of the consortia^33,34^, relying on strain co-dependence^32,44,45^, and being based on cell differentiation and thus being irreversible^36^. Furthermore, only one system has been shown to be effective for chemical production^33^; however, this relied on the drastic natural differences in growth rates between *E. coli* and yeast, allowing to control the composition by modulating the growth rate of only *E. coli* with light, while leaving the yeast unresponsive to light (“blind”). This strategy presents additional limitations for chemical production, because growth rates of microbial members are often similar, and because it does not allow for more than two-membered consortia.

Our multichromatic optogeneticTA system provides these missing capabilities. Blue light, red light, NIR light, and darkness afford orthogonal growth control over four *E. coli* strains simultaneously, dynamically regulating population ratios in co-cultures that would otherwise be easily destabilized by inherent growth-rate differences between members or stochastic effects of initial inoculations. A fifth, blind strain with a substantially lower growth rate than the bacterial strains could in principle be added to the co-cultures, with its abundance set indirectly through control of the other four. This unprecedented ability to not only stabilize but also dynamically control up to five-membered consortia, allowing for reversible shifts in microbial composition simply by changing illumination patterns on bioreactors brings engineered consortium-based chemical production appreciably closer to practical use.

Our systems offer the flexibility of controlling population compositions either by pulsing light patterns (alternating pulses of blue, red, and NIR, and darkness), or light intensities of different colors. We used modulated intensity to tune phenol-producing co-cultures because using modulated pulsing lowers the overall biomass at intermediate light setpoints (Figs. 2b and 3b) while using intensity modulation does not. We attribute this to altered regulatory dynamics of the circuits under pulsed versus continuous actuation. Under pulsing, cells alternate between fully permissive and fully repressive states, driving rapid swings in MazF expression; the repressive phase of each duty cycle may impose a substantial transient growth burden, particularly given the short half-life of the MazE antitoxin. By contrast, simultaneous red and NIR illumination is expected to hold the histidine kinase in an intermediately active state, producing steadier toxin/antitoxin levels and mitigating the growth penalty.

The dynamic control afforded by optogenetic regulation of consortia has significant effects on phenol production, and the conditions explored here merely touch the surface of its potential. Using light to transiently favor growth of the upstream strain promotes tyrosine accumulation before shifting light conditions mid-cultivation to favor the downstream strain to maximize tyrosine conversion to phenol. Although light-directed dynamic ratio control clearly enhances phenol production, our experiments sample only a small region of a very large, multidimensional and interdependent variable space. Other variables include the dialing of the red light intensity while keeping NIR constant or varying simultaneously. Future structured design-of-experiments approaches will map this space far more efficiently than the iterative optimization used here to approximate the true limits of this system. Moreover, feedback from fluorophore-labeled microbial consortia components and genetically encoded biosensors^12^ will enable closed-loop control, cybergenetic approaches, where the co-culture state is tracked in real time to maintain optimal conditions to maximize chemical production. The mathematical models presented here can support feedback control strategies, particularly model-based control. A promising future direction for such applications is their integration with data-driven components (e.g. machine-learning approaches), yielding hybrid models that improve predictive accuracy and may reduce the required mechanistic detail while preserving physical interpretability^46–48^. The ability to adjust light inputs instantly and reversibly in real time to optimize compositional trajectories and automate bioreactor operation could transform the use of microbial co-cultures for biomanufacturing.

Optogenetic control of microbial co-cultures offers several advantages over strategies based on cross-feeding or co-dependency. First, our optogenetic circuits are largely orthogonal to wild-type microbial metabolism. Cross-feeding or co-dependency strategies often require each strain to produce essential metabolites or nutrients for others, which can increase metabolic burden. In contrast, optogenetic regulation avoids the need for such additional metabolic load. Second, our multichromatic control over *mazEF* expression is orthogonal across strains, enabling independent regulation rather than complex interdependence between population members. Lastly, the ability to modulate light inputs with quick on–off switching enables real-time feedback control, unlike chemical inducers or metabolite-based strategies that lack precise temporal programmability and rapid responsiveness.

Co-cultures for phenol production serve as an effective proof of concept to demonstrate this technology, because of its relative simplicity and previous demonstration that co-cultures are more productive than monocultures^12^. However, this approach should be applicable to more complex biosynthetic pathways for higher-value products, such as pharmaceutical, agrochemicals, or nutraceuticals. These pathways are more likely to benefit from co-cultures involving more than two members^5,10,18,19^; in fact, microbial consortia may be the only feasible strategy to produce some of these products through cultivations.

While light penetration was not an impediment in this and other optogenetically controlled bioproduction platforms operating in lab-scale bioreactors^49–52^, translating these technologies to commercial scales will probably require additional process innovations and optimization. Mutations, gene circuits, and other genetic interventions have been shown to make microbial strains significantly more sensitive to light^51,53–56^, allowing for efficient optogenetic control in bioreactors of up to 5L at optical densities as high as OD_600_:∼300^51^. Additionally, light delivery will need to be enhanced in large scale bioreactors. Industrial or lab scale photobioreactors developed for photosynthetic organisms, such as microalgae and cyanobacteria^57,58^, may be adapted for optogenetically controlled bioprocesses. Other strategies include efficiently mixed bioreactors with submerged LEDs, thin reactor geometries, and external LED illumination systems through which cultures are recirculated^59,60^. The high light sensitivity of our system is an advantage in this context. The lower blue light dose (Fig. 2c) required at shorter forcing periods implies that less overall illumination is needed to tune the population while much more light-sensitive versions of the histidine kinases, including red- and NIR-responding ones, are available^61–63^. In addition, because dark-reversion in the RedTA and NIRTA systems is very slow, brief light pulses can be sufficient to drive consortia to a desired composition, and keep them there until the population is perturbed by natural drift or a new composition is desired. The ability to dynamically control microbial consortia, especially if closed-loop controls are implemented, could help realize the full potential of co-culture cultivations and apply them for the sustainable production of a wide range of molecules.

## Methods

### Strain construction and plasmid design

All plasmids and *Escherichia coli* strains used in this study are listed in Supplementary Tables 2 and 3, respectively. Plasmids were cloned via Gibson assembly and transformation into chemically competent DH5α strains, plated on LB agar plates with antibiotics. Antibiotic concentrations used in this work are as follows: 100 µg/mL carbenicillin, 50 µg/mL kanamycin, 100 µg/mL spectinomycin. Plasmids were purified from *Escherichia coli* using the Epoch DNA Miniprep.

The optogenetic toxin/antitoxin (optogeneticTA) and multichromatic superfolder GFP induction systems were constructed from pDawn, pRedawn, and pDmDERusk plasmids, gifts from Andreas Möglich (Addgene: #43796, #188978, and #213702, respectively), while H22P mutation was introduced by using site-directed mutagenesis on any pDawn-based plasmids. OptogeneticTA v2 and v3 versions were constructed by replacing kanamycin resistance gene with spectinomycin resistance gene in the optogenetic circuit backbone lacking or containing the *mazEF* circuit, respectively.

### Gene knockouts and genomic modifications

Deletion of endogenous *mazEF* genes was done using the Datsenko-Wanner method^64^. Linear DNA cassettes containing the kanamycin resistance marker were amplified from pKD4, flanked by 40 bp homology arms to the endogenous MazEF genes and electroporated into *E. coli* strains carrying the temperature-sensitive pKD46 plasmid. Following selection on kanamycin, the antibiotic marker was excised via the expression of FLP recombinase from the pCP20 plasmid, and knockouts were verified by colony PCR sequencing.

### Culture conditions and illumination setup

All cultures were grown in MY1 medium supplemented with 0.5% glucose and respective antibiotics at 37 °C and 200 rpm shaking. The recipe for MY1 media closely resembles the one reported by Guo et al. (2019)^12^. It is composed of 2 g/L NH_4_Cl, 5 g/L (NH_4_)_2_SO_4_, 3 g/L KH_2_PO_4_, 7.3 g/L K_2_HPO_4_, 8.4 g/L MOPS, 0.5 g/L NaCl, 0.24 g/L MgSO_4_, 340 mg/L thiamin, and 0.5 g/L yeast extract with trace elements. Trace elements contain 0.94 mg/L ZnCl_2_, 0.5 mg/L CoCl_2_, 0.38 mg/L CuCl_2_, 1.6 mg/L MnCl_2_, 3.77 mg/L CaCl_2_, and 3.6 mg/L FeCl_2_.

Monocultures were initiated from single colonies and grown overnight, then diluted to an optical density at 600 nm (OD₆₀₀) of 0.01 prior to illumination experiments in SensoPlate black 24-well glass-bottom or CytoOne 24-well plates. For accesing low blue-light intensities, we placed two neutral-density (ND9) filters beneath the 24-well plates, attenuating the blue light intensity emitted by the LEDs to ∼5.1% of the emitted intensity when measured. Cell growth was monitored by OD₆₀₀ measurements at various timepoints using a plate reader (Tecan Infinite M200PRO). sfGFP fluorescence intensity was measured using appropriate excitation/emission filters on Tecan plate reader (Infinite200PRO MPlex).

Illumination was done using OptoWells (OptoBiolabs) equipped with 447 nm (blue), 655 nm (red), and 775 nm (NIR) LEDs with light intensities ranging from 0-6.4, 0-3.3, and 0-2.83 mW/cm^2^, respectively. Pulsed and intensity-modulated illumination was controlled by custom OptoWELLControl v3.11.8 software, allowing independent control over different wavelengths at pulses and forcing periods ranging from 200 s to 8000 s. For experiments involving pulsing, the following intensities were used: blue light (0.32 mW/cm^2^), red light (0.17 mW/cm^2^), and NIR light (0.28 mW/cm^2^). For *E. coli* culturing, the OptoWells were mounted onto incubated shakers.

### Co-culture population analysis

For two-membered co-cultures, BlueTA and DarkTA strains were mixed 1:1 (a total OD_600_ 0.1 unless otherwise noted) and exposed to continuous or pulsed blue/dark light regimes using the OptoWELL instrument. RedTA/NIRTA co-cultures were analogously tested under red/NIR light inputs. Final population ratios were quantified after 24 hours for experiments in Fig. 3i and Supplementary Fig. 11-12 or 48 hours for all other experiments using a Beckman Coulter CytoFLEX S flow cytometer. Populations other than singlet cells were excluded based on the side scatter (SSC-A) vs forward scatter (FSC-A) plots. The four fluorescent proteins – superfolder GFP, mTagBFP2, mVenus, and mAmetrine1.2 – were gated following compensation of the four fluorophores with the following excitation laser and emission bandpass filter configurations: mTagBFP2 (excited at 405/10 nm, 450/45 nm filter), superfolder GFP (excited at 488/8 nm, 510/20 nm), mAmetrine1.2 (excited at 405/10 nm, 525/40 nm filter), and mVenus (excited at 488/8 nm, 550/30 nm filter). Pulsed experiments were conducted using forcing periods of 200 s, 2000 s, 4000 s, or 8000 s with varied duty cycles. Dynamic response experiments were performed by switching illumination programs mid-culture, as described in the Results section.

For three- and four-membered consortia, combinations of BlueTA, DarkTA, RedTA, and NIRTA were co-inoculated at equal or specified initial ratios to a total of OD 0.15 (three-membered) or 0.1 (four-membered). Blue/dark and red/NIR pulse programs were applied independently to achieve orthogonal control of subpopulation composition. Time-course sampling was performed every 1.5-3 hours to assess population ratios and total biomass using the flow cytometer.

### Phenol production experiments

For the upstream tyrosineproducing plasmid (pJJ426), aroA, aroG^fbr^, aroE, aroL, tyrA^fbr^ (feedback-resistant version), and aroC were cloned in a polycistronic manner under a constitutive proD promoter into a pETDuet-1 plasmid with lacI removed from the backbone. For the downstream phenol-producing plasmid (pJJ427), tyrosine-phenol lyase (tpl) enzyme from *Pasteurella multocida* was placed under the proD promoter in the pETDuet-1 plasmid lacking lacI. For the upstream strain, BL21·DE3 ΔtyrR (a gift from Prof. Hal Alper) was used to create a strain lacking *mazEF* and *recA* (BL21·DE3 ΔtyrRΔmazEFΔrecA, eJJ75). pJJ426 was transformed into eJJ75 to afford eJJ133. For the downstream strain, *mazEF* genes were knocked out of MG1655 ΔrecA ΔendA (DE3) (a gift from Prof. Kristala Prather; Addgene #37854) to afford eJJ67. pJJ427 was transformed into eJJ67 to generate the downstream phenol-producing strain, eJJ134.

For the NIRTA-controlled upstream strain, pJJ480 and pJJ470 were transformed into eJJ133 while pEY7 and an empty vector pJJ350 were transformed into the same strain for control. For the RedTA-controlled downstream strain, pJJ302 and pJJ383 were transformed into eJJ134 while pEY4 and pJJ350 were transformed into the same strain as control.

For the co-culture experiments, both upstream and downstream strains were cultured overnight in LB under permissive light conditions (NIR light for upstream and red light for downstream, under ambient light for control strains). The next day, upstream and downstream strains were mixed to a total OD of 0.2 in MY1 media at various inoculation ratios based on the counts per µL measured on the flow cytometer. Co-cultures were cultured in SensoPlate black 24 well glass bottom plates at 37 °C, 200 rpm for a total of 48 hours. If necessary, small aliquots of the cells (< 5 µL) were diluted into 1× PBS at different time points for population composition measurements on the flow cytometer. Following 48 hours, cells were harvested via centrifugation, and the supernatant was collected for HPLC analysis using a C18 column (ALLTIMA C18 5 MICRON, Cat. No. 88056, Avantor). The column was maintained at 35 °C throughout the run with 0.1% formic acid in water or acetonitrile as the mobile phase with the following sequence of acetonitrile percentage: 10-30% for 17 minutes, 30-50% for 3 minutes, 50-100% for 0.5 minutes, 100% for 2.5 minutes, 100-10% for 0.5 minutes, 10% for 5.5 minutes. Tyrosine was detected using a DAD detector at 230 nm while phenol was detected using 280 nm.

### Model fitting and simulations

Parameter estimation was performed with COPASI^65^, minimizing a weighted sum of squared errors against experimental biomass time-course data. Unmeasured initial conditions were estimated jointly with model parameters; for consortium fits, single-strain estimates served as initial guesses with relaxed bounds. For simplicity, duty cycles in pulse-width-modulated experiments were normalized to a continuous input between 0 and 1, corresponding to 0% and 100% duty cycle, respectively. Numerical integration was performed using COPASI’s default deterministic time-course method, which employs LSODA for standard simulations and LSODAR when event detection is required. Fitted models are provided as Systems Biology Markup Language (SBML) files at https://github.com/sespirio/Models_optogeneticTA. Additional details are provided in Supplementary Discussion 1 and Supplementary Table 1.

### Statistics

Experiments were performed with at least three biological replicates. Comparison between two groups were analyzed by unpaired t-tests with significance level at P<0.05 using GraphPad Prism 10.6.1. All data are shown as mean and error bars from standard deviation.

## Acknowledgements

This work was supported by the U.S. Department of Energy, Office of Biological and Environmental Research (J.L.A. – Award Number DE-SC0022155) and U.S. National Science Foundation (MCB-2300239). J.J. was supported by the Omenn-Darling Postdoctoral Fellowship.

## Author Contributions

J.J. and J.L.A. conceived the study and designed the experiments. J.J. designed and constructed the optogenetic toxin-antitoxin constructs/strains and performed experimental characterizations. E.Y. and T.N.S. contributed to plasmid construction and assisted with flow cytometry measurements. J.J. designed and performed the phenol production experiments and analysis. E.A. developed the mathematical framework. S.E.-R. performed parameter estimation and model validation. J.J., E.A., and S.E.-R. analysed the data. J.L.A. supervised the project and acquired funding. J.J., E.A., S.E.-R., and J.L.A. wrote the manuscript and created figures. All authors edited and approved the manuscript.

## Data and Code Availability

Plasmids and strains in this study are available upon request. Fitted models are provided as Systems Biology Markup Language (SBML) files on GitHub.

