## Supplementary Information for "Multichromatic Dynamic Control of Multi-Membered Microbial Consortia Compositions for Chemical Production"

### Supplementary Discussion 1

#### Modeling

Here, we present a modular mathematical framework to quantitatively describe the behavior of our light-controllable engineered strains. We employ commonly used assumptions regarding gene expression and cell growth kinetics<sup>1,2</sup>. Furthermore, the main modeling principles for the interactions involving *YFI*, *FixJ* and *mazE*, *mazF* are based on Hennemann *et al.* (2018)<sup>3</sup> and Gelens *et al.* (2013)<sup>4</sup>, respectively.

The behavior of a single bacterial strain (monoculture) can be described by the following Ordinary Differential Equation (ODE) system:

$$\begin{aligned} \frac{d[Y_{DD}]}{dt} &= b_Y g_1([F]) - 2g_{2i}(I_B, I_R, I_{IR})[Y_{DD}] \\ &\quad + g_{3i}(I_B, I_R, I_{IR})([Y_{LD}] + [Y_{DL}]) - \mu([N], [F])[Y_{DD}] \end{aligned} \quad (1a)$$

$$\begin{aligned} \frac{d[Y_{LD}]}{dt} &= -(g_{3i}(I_B, I_R, I_{IR}) + g_{2i}(I_B, I_R, I_{IR}))[Y_{LD}] \\ &\quad + g_{2i}(I_B, I_R, I_{IR})[Y_{DD}] + g_{3i}(I_B, I_R, I_{IR})[Y_{LL}] - \mu([N], [F])[Y_{LD}] \end{aligned} \quad (1b)$$

$$\begin{aligned} \frac{d[Y_{DL}]}{dt} &= -(g_{3i}(I_B, I_R, I_{IR}) + g_{2i}(I_B, I_R, I_{IR}))[Y_{DL}] \\ &\quad + g_{2i}(I_B, I_R, I_{IR})[Y_{DD}] + g_{3i}(I_B, I_R, I_{IR})[Y_{LL}] - \mu([N], [F])[Y_{DL}] \end{aligned} \quad (1c)$$

$$\frac{d[Y_{LL}]}{dt} = -2g_{3i}(I_B, I_R, I_{IR})[Y_{LL}] + g_{2i}(I_B, I_R, I_{IR})([Y_{LD}] + [Y_{DL}]) - \mu([N], [F])[Y_{LL}] \quad (1d)$$

$$\begin{aligned} \frac{d[J]}{dt} &= b_J g_1([F]) - k_K[Y_{DD}][J] + k_H[J_P] \\ &\quad + k_P([Y_{LD}] + [Y_{DL}] + [Y_{LL}])[J_P] - \mu([N], [F])[J] \end{aligned} \quad (1e)$$

$$\begin{aligned} \frac{d[J_P]}{dt} &= k_K[Y_{DD}][J] - k_H[J_P] - k_P([Y_{LD}] + [Y_{DL}] + [Y_{LL}])[J_P] \\ &\quad - \mu([N], [F])[J_P] \end{aligned} \quad (1f)$$

$$\frac{d[C]}{dt} = g_4(J_P)g_1([F]) - (\gamma_C + \mu([N], [F]))[C] \quad (1g)$$

$$\frac{d[F]}{dt} = g_4(J_P)g_1([F]) - \mu([N], [F])[F] - 2\alpha_{TH}[F][E] \quad (1h)$$

$$\frac{d[E]}{dt} = g_5([C])g_1([F]) - (\gamma_E + \mu([N], [F]))[E] - \alpha_{TH}[F][E] \quad (1i)$$

$$\frac{d[P]}{dt} = \mu([N], [F])[P] \quad (1j)$$

$$\frac{d[N]}{dt} = -\delta\mu([N], [F])[P] \quad (1k)$$

where $i \in \{1, 2, 3, 4\}$, following 1 for *BlueTA*, 2 for *DarkTA*, 3 for *RedTA*, and 4 for *NIRTA* strains. Furthermore:

$$\begin{aligned} g_1([F]) &= \frac{K_F}{[F]^{n_F} + K_F}, \\ g_{2i}(I_B, I_R, I_{IR}) &= \begin{cases} g_{21}(I_B, I_R, I_{IR}) = k_I I_B, & \text{if } i = 1 \\ g_{22}(I_B, I_R, I_{IR}) = k_I, & \text{if } i = 2 \\ g_{23}(I_B, I_R, I_{IR}) = k_I I_R, & \text{if } i = 3 \\ g_{24}(I_B, I_R, I_{IR}) = k_I I_{IR}, & \text{if } i = 4, \end{cases} \\ g_{3i}(I_B, I_R, I_{IR}) &= \begin{cases} g_{31}(I_B, I_R, I_{IR}) = k_R, & \text{if } i = 1 \\ g_{32}(I_B, I_R, I_{IR}) = k_R I_B, & \text{if } i = 2 \\ g_{33}(I_B, I_R, I_{IR}) = k_R(I_{IR} + b_{D_1}), & \text{if } i = 3 \\ g_{34}(I_B, I_R, I_{IR}) = k_R(I_R + b_{D_2}), & \text{if } i = 4, \end{cases} \\ g_4([P]) &= k_J \frac{[P]^{n_J}}{[P]^{n_J} + K_J}, \\ g_5([C]) &= k_C \frac{K_C}{[C]^{n_C} + K_C}, \\ \mu([N], [F]) &= g_1([F]) k_N \frac{N^{n_g}}{N^{n_g} + K_N} \end{aligned}$$

Note that $\mathbf{x} := [Y_{DD}, Y_{LD}, Y_{DL}, Y_{LL}, J, J_P, C, F, E, P, N]^T$ is the state vector. Thus, $\mathbf{x}(t_0) = \mathbf{x}_0$ represents the initial condition of the system. The control input is denoted as $u \in \{I_B, I_R, I_{IR}\}$, selected according to the strain. The parameter vector is defined as $\boldsymbol{\theta} := [b_Y, b_J, k_K, k_H, k_P, \gamma_C, \gamma_E, \gamma_P, \alpha_{TH}, \delta, K_F, k_I, k_R, k_J, K_J, k_C, K_C, k_N, K_N]^T$. The meaning of these variables, functions, and parameters is given in Supplementary Table 1.

To capture the dynamics of a microbial community of an arbitrary number of bacterial strains, we can use Equations (1a)-(1k) for each strain involved and replace Equation (1k) with:

$$\frac{d[N]}{dt} = -\sum_{j \in \mathcal{P}} \delta_j \mu_j([N], [F])[P_j],$$

where $\mathcal{P} \subseteq \{1, 2, 3, 4\}$ is the set of members in the consortium. Here, each strain contributes to the uptake dynamics of the shared nutrient. In the general case, model parameters and initial conditions (except for the shared initial nutrient concentration) are assumed specific to each strain in the consortium. Thus,

$$\boldsymbol{\theta} := [\boldsymbol{\theta}_j^T, \forall j \in \mathcal{P}]^T, \text{ and the state vector is redefined as: } \mathbf{x} := [N, \mathbf{x}_j^T, \forall j \in \mathcal{P}]^T,$$

$$\text{where } \mathbf{x}_j := [Y_{DD}]_j, [Y_{LD}]_j, [Y_{DL}]_j, [Y_{LL}]_j, [J]_j, [J_P]_j, [C]_j, [F]_j, [E]_j, [P]_j]^T.$$

##### Parameter Estimation

Parameter fitting of experiments involving single strains and two- and three-strain consortia was performed using the BasiCO Python library<sup>5</sup>, which interfaces with COPASI (Complex Pathway Simulator)<sup>6</sup>.

The loss function, $\mathcal{L}$, minimized during parameter optimization was a weighted sum of squared errors:

$$\mathcal{L} = \sum_{i,j} \omega_j (x_{i,j} - y_{i,j})^2,$$

where $x_{i,j}$ is an experimental data point and $y_{i,j}$ is the corresponding model prediction. The indices $i$ and $j$ denote rows (sampling times) and columns (measured states), respectively, while $\omega_j$ is the weight associated with measured state $j$. The weights were calculated from the mean of the squared values in each column. Unless otherwise stated, when dealing with the same biological system (single strain or consortia) under the same experimental and control conditions but with varying optogenetic manipulations, we fitted all experiments simultaneously, so that the loss function accounted for all experiments at once.

In general, we followed an *ensemble* parameter estimation strategy, where we first employed derivative-free global optimization algorithms to identify candidate parameter sets. These included two evolutionary algorithms, evolutionary programming<sup>7</sup> and a genetic algorithm<sup>8</sup>, based on the concept of populations and selection, and one stochastic algorithm, particle swarm optimization<sup>9</sup>, based on position and velocity concepts. Each algorithm was randomly initialized twice. Starting from the three best parameter sets obtained in this way, we then applied a direct search algorithm based on geometric concepts, the Hooke-Jeeves method<sup>10</sup>, which is a local optimization method. The final parameter set was selected as the best solution from this local search. Hyperparameters for all optimization methods were kept at their default values in COPASI<sup>6</sup>.

For simplicity, when fitting systems actuated with pulse-width modulation, pulsing cycles were normalized between 0 and 1 and treated as a continuous input in the model. In experiments actuated by intensity modulation, the light intensity was used directly as the input to the model.

*Remark on single-strain systems:* the initial conditions for biomass and substrate (Eqs. (1j), (1k)) were set according to the experimental measurements, whereas the remaining (unmeasured) initial conditions were estimated together with the model parameters. Since the inocula in the experiments were taken from non-growth-repressing conditions, the initial conditions for the intracellular states in Eqs. (1a)–(1g) were constrained around the non-repressing steady-state values predicted by the submodel in Hennemann et al (2018)<sup>3</sup>.

To establish biologically sound ranges for the unknown model parameters in Eqs. (1a)–(1g), we performed a set of preliminary fits with simulated data using parameter values from Hennemann et al (2018)<sup>3</sup>, extending the model to account for production and growth/dilution effects neglected in the original formulation, and assuming a constant maximum growth rate (estimated from experiments) with no toxin–antitoxin dynamics. The resulting parameter values were used as a basis to constrain the parameter estimation, complemented by literature-based initial guesses (such as for $\gamma_{ME}$<sup>11</sup> and for $\alpha_{TH}$, initially chosen such that $\alpha_{TH} \gg \mu^4$). In general, these parameter values (either estimated from submodels as mentioned above or taken from the literature) were assigned bounds that were sufficiently wide to facilitate convergence and loss minimization. Additionally, the bounds for the maximum growth rate and the corresponding substrate yields (assuming full substrate consumption under non-repressive conditions) were set directly from experiments.

*Remarks on consortia systems:* based on the parameter sets obtained for the individual optogenetic strains, we used those parameters as initial guesses for the corresponding strains in the consortia models and relaxed their bounds to be sufficiently wide to facilitate convergence of the minimization, allowing for additional uncertainty introduced by inter-strain interactions. As before, the bounds for the maximum growth rate and the corresponding substrate yields for each strain were set directly from the relevant consortium experiments, assuming full substrate consumption under non-repressive conditions and no noticeable growth of competitors.

All fitted models are shared as SBML files, in which all model equations and fitted parameter values are available. It should be noted that, while the model structure (states and interactions) is derived from mechanistic knowledge of the system, only biomass concentrations were measured over time and all remaining states are unobserved. Consequently, multiple parameter sets can in principle fit the experimental data, and the model is therefore not uniquely identifiable. For this reason, the model should be viewed primarily as a mechanistically grounded structural representation that is consistent with the observed macroscopic behavior, rather than as a precise quantitative estimate of the underlying parameter values. This lack of uniqueness is expected given the large number of states and parameters relative to the available measurements. The reported parameter sets should thus be interpreted as representative solutions consistent with the data under the imposed constraints.

| Symbolism | Description |
| --- | --- |
| $[Y_{DD}], [Y_{LD}], [Y_{DL}], [Y_{LL}]$ | Concentrations of *YFI*, with its two *LOV* photosensors, corresponding to the subscript-designated states: <sub>D</sub> (dark-adapted) and <sub>L</sub> (light-adapted) |
| $[J], [J_P]$ | Concentrations of *FixJ* in dephosphorylated (denoted without a subscript) and phosphorylated (denoted with the subscript <sub>p</sub>) states |
| $[C]$ | Concentration of *cI* |
| $[F]$ | Concentration of *mazF* |
| $[E]$ | Concentration of *mazE* |
| $[P]$ | Concentration of a bacterial strain |
| $[N]$ | Concentration of the nutrient |
| $I_B, I_R, I_{IR}$ | Applied intensities of blue, red, infrared light, respectively |
| $b_Y, b_J,$ | Maximal expression rates with respect to $[Y_{DD}], [J]$, respectively |
| $k_K$ | Phosphorylation rate constant of *FixJ* |
| $k_P$ | Dephosphorylation rate constant of *FixJ* |
| $k_H$ | Rate constant of spontaneous hydrolysis of the phosphoryl group on *FixJ* |
| $\gamma_C, \gamma_E$ | Degradation rate constants with respect to *cI*, *mazE* respectively |
| $\alpha_{TH}$ | Binding rate constant of *mazF* and *mazE* |
| $\delta$ | Yield parameter defined as the inverse of number of bacteria generated from the nutrient concentration |
| $g_1([F])$ | Hill function describing the inhibition of cell growth induced by *mazF* |
| $g_{2i}(I_B, I_R, I_{IR})$ | Function representing the photoactivation rate constant of *LOV* photosensors with respect to *YF1* |
| $g_{3i}(I_B, I_R, I_{IR})$ | Function representing the dark recovery rate constant of *LOV* photosensors with respect to *YF1* |
| $g_4([J_P])$ | Hill function describing the activation of *cI* expression by (phospho-)*FixJ* |
| $g_5([C])$ | Hill function describing the repression of *mazE* expression by *cI* |
| $\mu([N], [F])$ | Function describing cell growth - it is defined as the multiplication of $g_1([F])$ with a repression Hill function related to the nutrient concentration |

**Supplementary Table 1.** List of notations and their meanings for the introduced mathematical framework with respect to a strain *i*. Suffix denoting specific bacterial strains: 1 for *BlueTA*, 2 for *DarkTA*, 3 for *RedTA*, and 4 for *NIRTA*.

| Plasmid Name | Content | Antibiotic Marker |
| --- | --- | --- |
| pET24b |  | Kanamycin |
| pKD4 | See Datsenko & Wanner (2000)<sup>12</sup> | Kanamycin and Carbenicillin |
| pKD46 | See Datsenko & Wanner (2000)<sup>12</sup> | Carbenicillin |
| pCP20 | See Datsenko & Wanner (2000)<sup>12</sup> | Carbenicillin |
| pJJ91 | lacI<sup>q</sup> (HO1_DrBPhP_FixJ) pFixK2 *cI* pR_sfGFP | Kanamycin |
| pJJ502 | lacI<sup>q</sup> (YF1_FixJ) pFixK2 *cI* pR_sfGFP | Kanamycin |
| pJJ475 | lacI<sup>q</sup> (YF1<sup>H22P</sup>_FixJ) pFixK2 *cI* pR_sfGFP | Kanamycin |
| pJJ412 | lacI<sup>q</sup> (HO1_DmF1_FixJ) pFixK2 *cI* pR_sfGFP | Kanamycin |
| pJJ331 | J23100_sfGFP | Kanamycin |
| pJJ426 | proD tyrA<sup>fbr</sup> AroG<sup>fbr</sup> AroE AroL AroA AroC | Carbenicillin |
| pJJ427 | Prod_tpl | Carbenicillin |
| pJJ350 | Empty vector *cloDF13 ori* | Spectinomycin |
| pJJ270 | lacI<sup>q</sup> (YF1_FixJ) pFixK2 *cI* pFixK2_MazF_pR_MazE | Kanamycin |
| pJJ483 | lacI<sup>q</sup>_(YF1<sup>H22P</sup>_FixJ)_pFixK2_*cI*_pFixK2_MazF_pR_MazE | Kanamycin |
| pJJ186 | lacI<sup>q</sup>_(HO1_DrBPhP_FixJ)_pFixK2_*cI*_pFixK2_MazF_pR_MazE | Kanamycin |
| pJJ460 | lacI<sup>q</sup>_(HO1_DmF1_FixJ)_pFixK2_*cI*_pFixK2_MazF_pR_MazE | Kanamycin |
| pJJ310 | lacI<sup>q</sup>_(YF1_FixJ)_pFixK2_*cI*_pFixK2_MazF_pR_MazE_J23109_mTagBFP2 | Kanamycin |
| pJJ492 | lacI<sup>q</sup>_(YF1<sup>H22P</sup>_FixJ)_pFixK2_*cI*_pFixK2_MazF_pR_MazE_J23109_sfGFP | Kanamycin |
| pJJ268 | lacI<sup>q</sup>_(HO1_DrBPhP_FixJ)_pFixK2_*cI*_pFixK2_MazF_pR_MazE_J23109_mAmetrine1.2 | Kanamycin |
| pJJ480 | lacI<sup>q</sup>_(HO1_DmF1_FixJ)_pFixK2_*cI*_pFixK2_MazF_pR_MazE_J23109_mVenus | Kanamycin |
| pJJ379 | lacI<sup>q</sup>_(YF1_FixJ)_pFixK2_*cI*_pR_MCS | Spectinomycin |
| pJJ473 | lacI<sup>q</sup>_(YF1<sup>H22P</sup>_FixJ)_pFixK2_*cI*_pR_MCS | Spectinomycin |
| pREDawn-StrR-MCS (pJJ61) | lacI<sup>q</sup>_(HO1_DrBPhP_FixJ)_pFixK2_*cI*_pR_MCS<br>See Multamäki et al. (2022)<sup>13</sup> | Spectinomycin |
| pJJ470 | lacI<sup>q</sup>_(HO1_DmF1_FixJ)_pFixK2_*cI*_pR_MCS | Spectinomycin |
| pJJ385 | lacI<sup>q</sup>_(YF1_FixJ)_pFixK2_*cI*_pFixK2_MazF_pR_MazE | Spectinomycin |
| pJJ295 | lacI<sup>q</sup>_(YF1<sup>H22P</sup>_FixJ)_pFixK2_*cI*_pFixK2_MazF_pR_MazE | Spectinomycin |
| pJJ383 | lacI<sup>q</sup>_(HO1_DrBPhP_FixJ)_pFixK2_*cI*_pFixK2_MazF_pR_MazE | Spectinomycin |
| pJJ464 | lacI<sup>q</sup>_(HO1_DmF1_FixJ)_pFixK2_*cI*_pFixK2_MazF_pR_MazE | Spectinomycin |
| pEY4 | lacI<sup>q</sup>_(YF1_FixJ)_pFixK2_*cI*_pR_MCS_J23109_mTagBFP2 | Kanamycin |
| pEY5 | lacI<sup>q</sup>_(YF1_FixJ)_pFixK2_*cI*_pR_MCS_J23109_sfGFP | Kanamycin |
| pEY6 | lacI<sup>q</sup>_(YF1_FixJ)_pFixK2_*cI*_pR_MCS_J23109_mAmetrine1.2 | Kanamycin |
| pEY7 | lacI<sup>q</sup>_(YF1_FixJ)_pFixK2_*cI*_pR_MCS_J23109_mVenus | Kanamycin |
| pJJ302 | lacI<sup>q</sup>_(HO1_DrBPhP_FixJ)_pFixK2_*cI*_pFixK2_MazF_pR_MazE_J23109_mTagBFP2 | Kanamycin |

**Supplementary Table 2.** Plasmids constructed and used in this work. Brackets indicate polycistronic operons.

| Strain Name | Content | Source |
| --- | --- | --- |
| MG1655·DE3 ΔendA ΔrecA |  | Addgene #37854 |
| BL21·DE3 ΔtyrR |  | Brooks et al. (2023)<sup>14</sup> |
| eJJ75 | BL21·DE3 ΔtyrR ΔmazEF ΔrecA | This work |
| eJJ67 | MG1655·DE3 ΔendA ΔrecA ΔmazEF | This work |
| eJJ10 | eJJ67 + pJJ502 | This work |
| eJJ11 | eJJ67 + pJJ475 | This work |
| eJJ12 | eJJ67 + pJJ91 | This work |
| eJJ13 | eJJ67 + pJJ412 | This work |
| eJJ14 | eJJ67 + pJJ331 | This work |
| eJJ7 | eJJ67 + pJJ270 | This work |
| eJJ8 | eJJ67 + pJJ483 | This work |
| eJJ6 | eJJ67 + pJJ186 | This work |
| eJJ24 | eJJ67 + pJJ460 | This work |
| eJJ81 | eJJ67 + pJJ270 + pJJ350 | This work |
| eJJ82 | eJJ67 + pJJ483 + pJJ350 | This work |
| eJJ83 | eJJ67 + pJJ186 + pJJ350 | This work |
| eJJ84 | eJJ67 + pJJ460 + pJJ350 | This work |
| eJJ85 | eJJ67 + pJJ270 + pJJ379 | This work |
| eJJ99 | eJJ67 + pJJ483 + pJJ473 | This work |
| eJJ100 | eJJ67 + pJJ186 + pJJ61 | This work |
| eJJ101 | eJJ67 + pJJ460 + pJJ470 | This work |
| eJJ15 | eJJ67 + pJJ270 + pJJ385 | This work |
| eJJ16 | eJJ67 + pJJ483 + pJJ295 | This work |
| eJJ17 | eJJ67 + pJJ186 + pJJ383 | This work |
| eJJ18 | eJJ67 + pJJ460 + pJJ464 | This work |
| eJJ19 | eJJ67 + pET24b | This work |
| eJJ20 | eJJ67 + pET24b + pJJ350 | This work |
| eJJ150 | eJJ67 + pJJ310 + pJJ385 | This work |
| eJJ151 | eJJ67 + pJJ492 + pJJ295 | This work |
| eJJ152 | eJJ67 + pJJ268 + pJJ383 | This work |
| eJJ153 | eJJ67 + pJJ480 + pJJ470 | This work |
| eJJ154 | eJJ67 + pEY4 + pJJ350 | This work |
| eJJ155 | eJJ67 + pEY5 + pJJ350 | This work |
| eJJ156 | eJJ67 + pEY6 + pJJ350 | This work |
| eJJ157 | eJJ67 + pEY7 + pJJ350 | This work |
| eJJ133 | eJJ75 + pJJ426 | This work |
| eJJ134 | eJJ67 + pJJ427 | This work |
| eJJ173 | eJJ133 + pEY7 + pJJ350 | This work |
| eJJ174 | eJJ133 + pJJ480 + pJJ470 | This work |
| eJJ175 | eJJ134 + pEY4 + pJJ350 | This work |
| eJJ176 | eJJ134 + pJJ302 + pJJ383 | This work |

**Supplementary Table 3.** Strain list used in this work

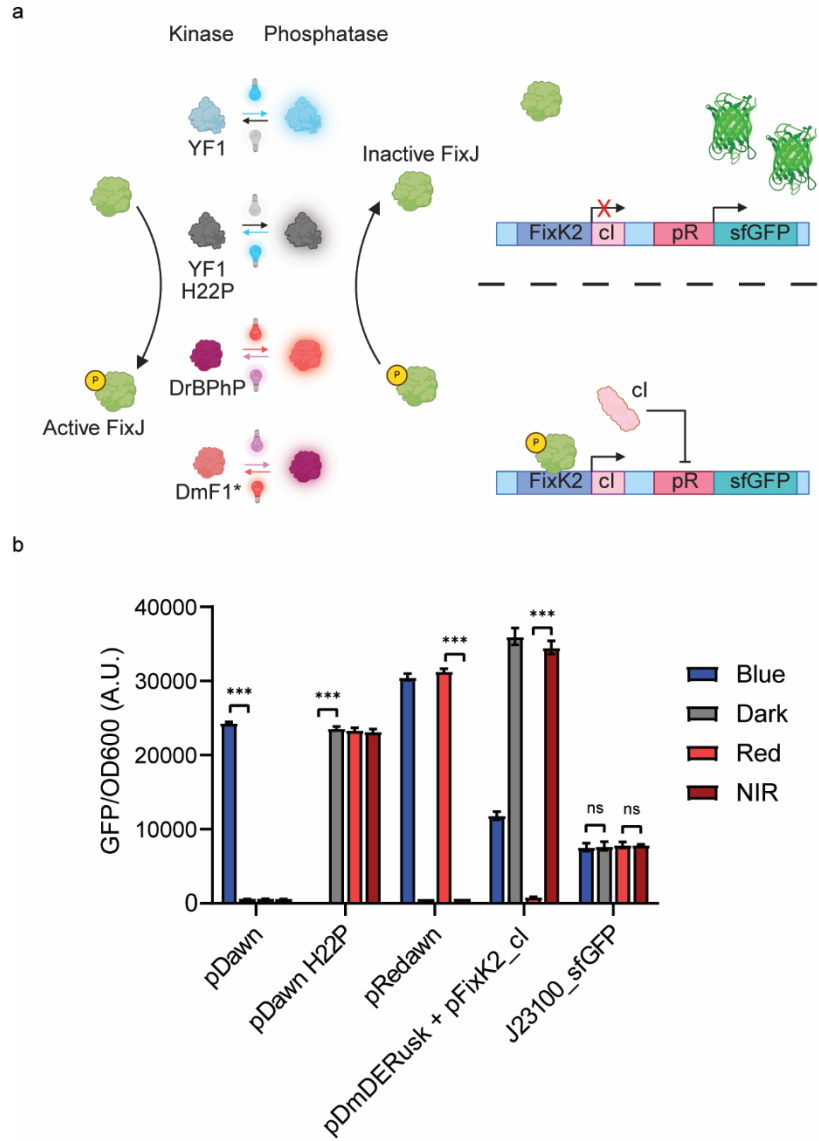

**Supplementary Figure 1.** Applying the YF1<sup>H22P</sup> mutant to the pDawn, pRedawn, and a modified pDmDERusk system. **a** Schematic of the light-inducible gene expression system (superfolder GFP reporter) using four variants of the photoresponsive histidine kinases, YF1, YF1<sup>H22P</sup>, DrBPhP, DmF1<sup>3,13,15,16</sup>. **b** For the pDawn system, we observed ~43-fold higher superfolder GFP expression under blue light over darkness while the YF1<sup>H22P</sup> mutant version yielded ~163-fold higher expression in darkness over blue light, an inverted response to the pDawn system.<sup>15</sup> The pREDawn system<sup>13</sup>, which involves a red light-activated phosphatase – *DrBPhP* – in place of YF1 in pDawn, demonstrated a red light-inducible and NIR-repressible gene expression system up to 43-fold. Lastly, we utilized the NIR-activated phosphatase, DmF1 mutant, from the pDmDERusk system<sup>16</sup> in place of *DrBPhP*, allowing genes to be induced with near-infrared light at 39-fold over red light. Statistics are derived from t-test (\*\*\*) $P < 0.001$,

\*\*P<0.002, \*P<0.033, ns: not significant). Blue, red, and NIR light were irradiated with intensities of 0.32, 0.17, and 0.28 mW/cm<sup>2</sup>, respectively. All data are shown as mean values with error bars representing standard deviation from n=3 biological replicates.

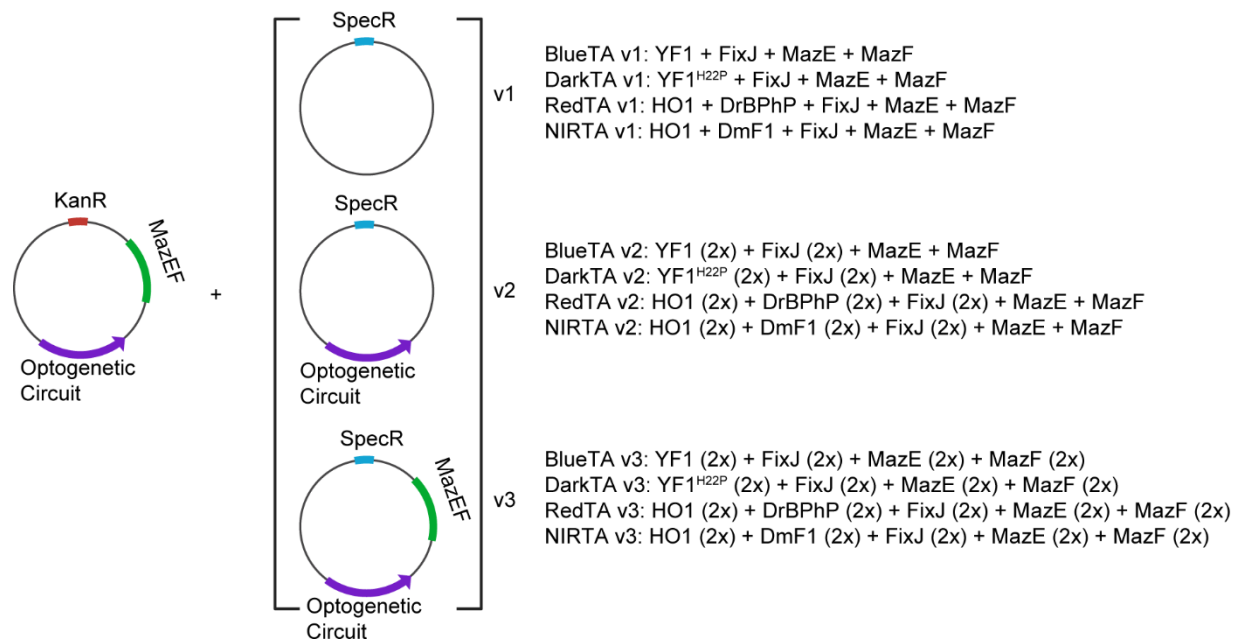

**Supplementary Figure 2.** Schematic overview of the plasmid-based circuit designs used to construct light-responsive toxin-antitoxin systems (optogeneticTA). The first plasmid backbone encodes photosensory components together with the *mazEF* toxin-antitoxin. Three design iterations (v1-v3) were generated to systematically vary circuit performance. v1 contains an additional empty SpecR plasmid to match the antibiotic resistances of v2 and v3. v2 contains an extra copy of photosensory components (without the additional toxin/antitoxin). v3 contains an extra copy of photosensory components as well as the toxin/antitoxin pair.

**a** *BlueTA monoculture model fitting at various blue/dark pulse duty cycles ($t_{FP} = 2000$ s)*

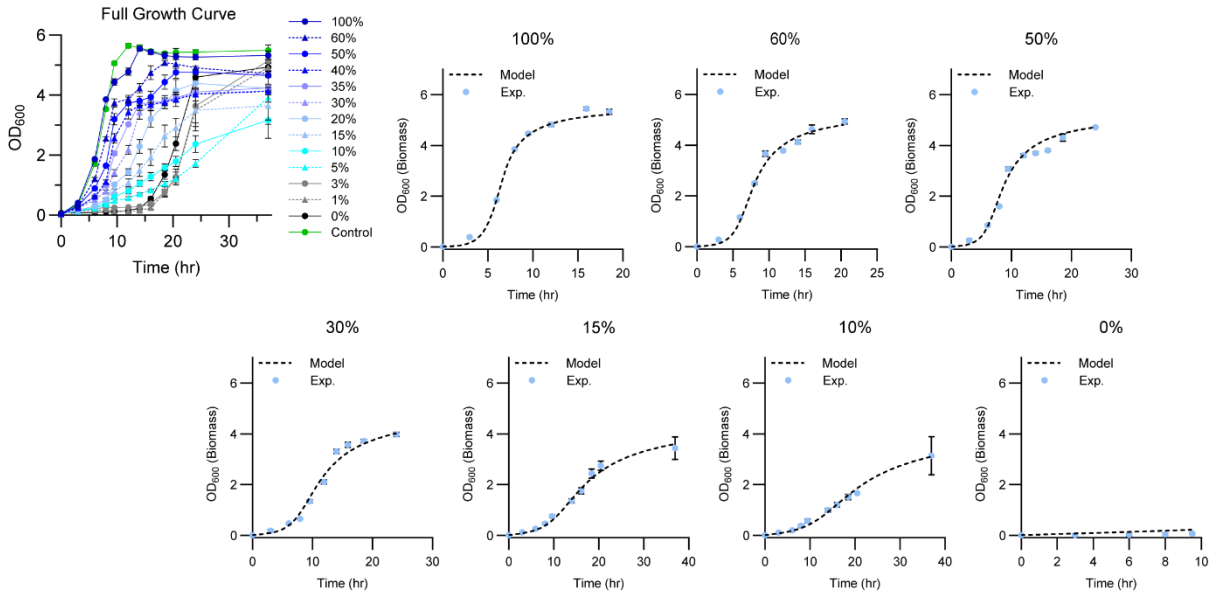

**b** *DarkTA monoculture model fitting at various blue/dark pulse duty cycles ($t_{FP} = 2000$ s)*

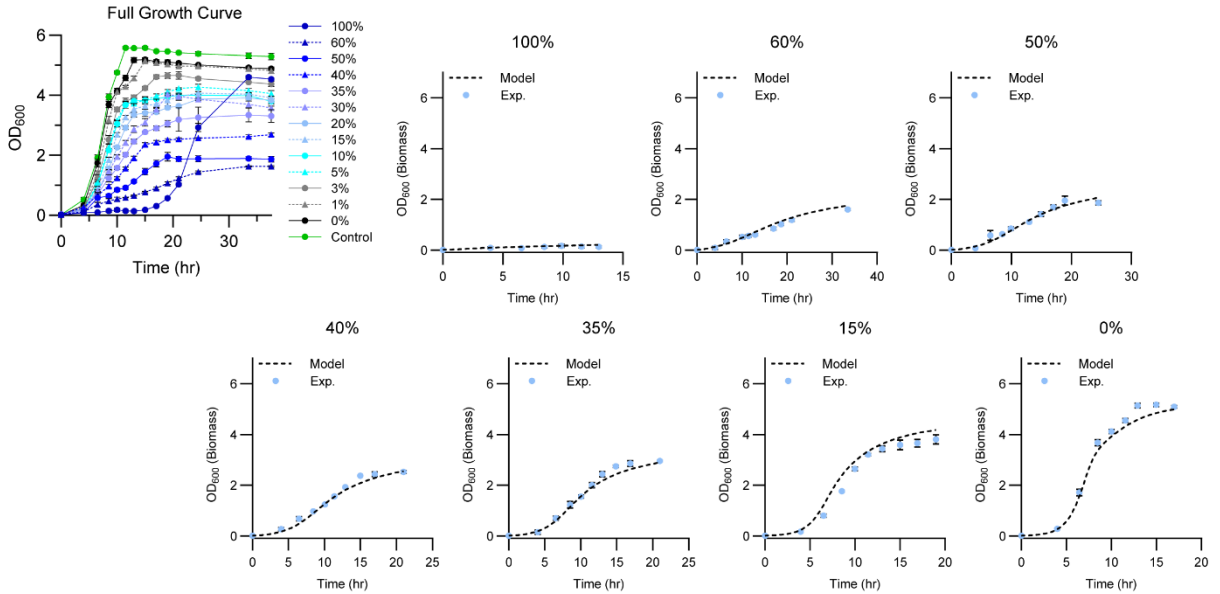

**Supplementary Figure 3.** Full growth curve time courses of BlueTA (a) and DarkTA (b) monocultures across blue light/dark pulsing cycles (at $0.32 \text{ mW/cm}^2$ of blue light) and the mechanistic model fit. **a** Fits of BlueTA monoculture growth model to experimental biomass trajectories under varying blue light pulsing cycles (100, 60, 50, 30, 15, 10, and 0% at $t_{FP}=2000$ s). n% of blue/dark pulse indicates n% of blue light irradiation, followed by (100-n)% of darkness at a certain $t_{FP}$. **b** Corresponding model fits for DarkTA monoculture under the indicated blue-light pulsing cycles (100, 60, 50, 40, 35, 15, and 0%). In all panels, biomass was measured as optical density at 600 nm ($OD_{600}$) at indicated hours. Blue circles denote

experimental measurements, and dotted lines indicate model predictions. Model fitted data are shown as mean values with error bars representing standard deviation from n=3 biological replicates while the full growth curve is from n=4 biological replicates.

**a** *RedTA monoculture model fitting at various red/NIR pulse duty cycles ($t_{FP} = 2000$ s)*

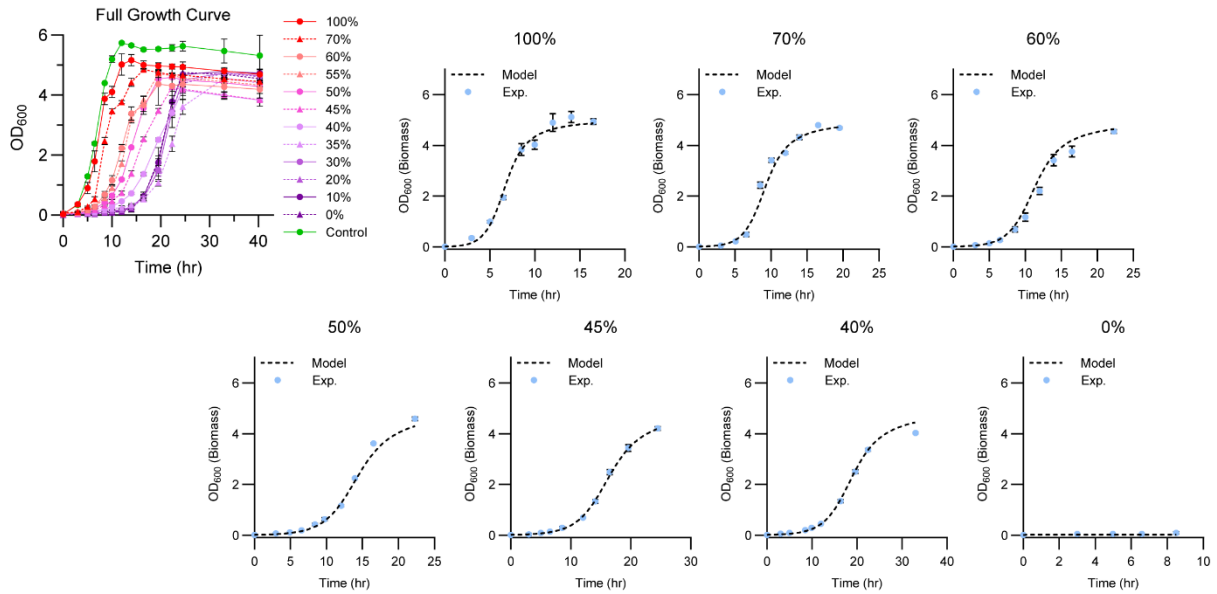

**b** *NIRTA monoculture model fitting at various red/NIR pulse duty cycles ($t_{FP} = 2000$ s)*

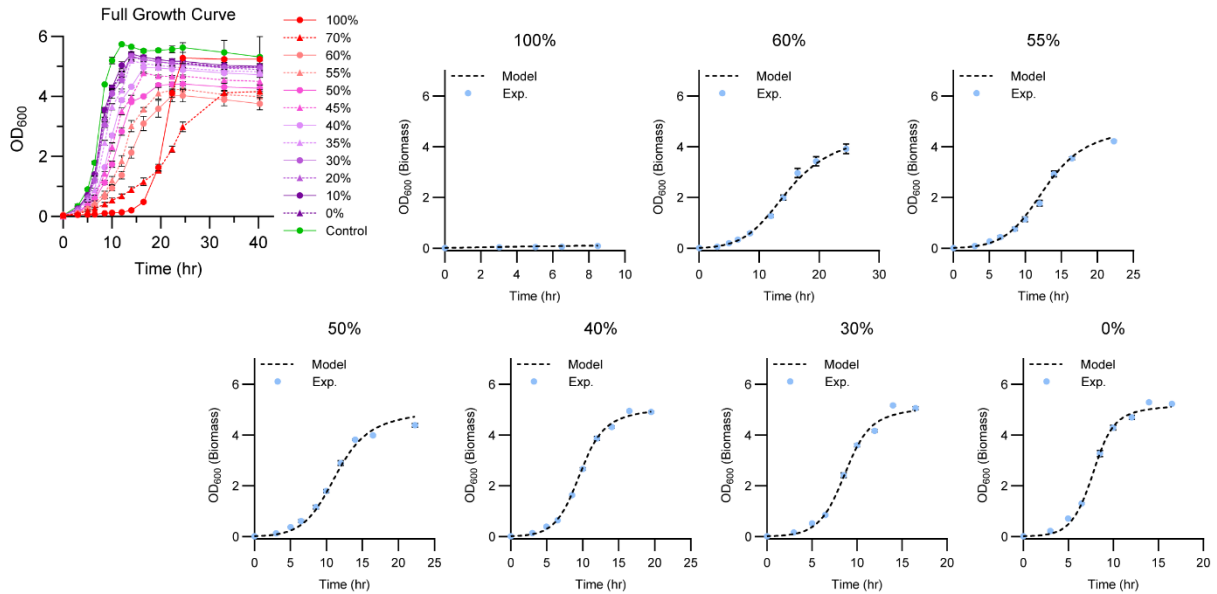

**Supplementary Figure 4.** Full growth curve time courses of RedTA (a) and NIRTA (b) monocultures across red/NIR light pulsing cycles (at 0.17 and 0.28 mW/cm<sup>2</sup>, respectively) and the mechanistic model fit. n% of Red/NIR pulse indicates n% of red light irradiation, followed by (100-n)% of NIR light irradiation at a certain $t_{FP}$. **a** Fits of RedTA monoculture growth model to experimental biomass trajectories under varying red/NIR light pulsing cycles (100, 70, 60, 50,

45, 40, and 0% at $t_{FP}=2000$ s). **b** Corresponding model fits for NIRTA monoculture under the indicated red/NIR light pulsing cycles (100, 60, 55, 50, 40, 30, and 0%). In all panels, biomass was measured as optical density at 600 nm ($OD_{600}$) at indicated hours. Blue circles denote experimental measurements, and dotted lines indicate model predictions. Model fitted data are shown as mean values with error bars representing standard deviation from $n=3$ biological replicates while the full growth curve is from $n=4$ biological replicates.

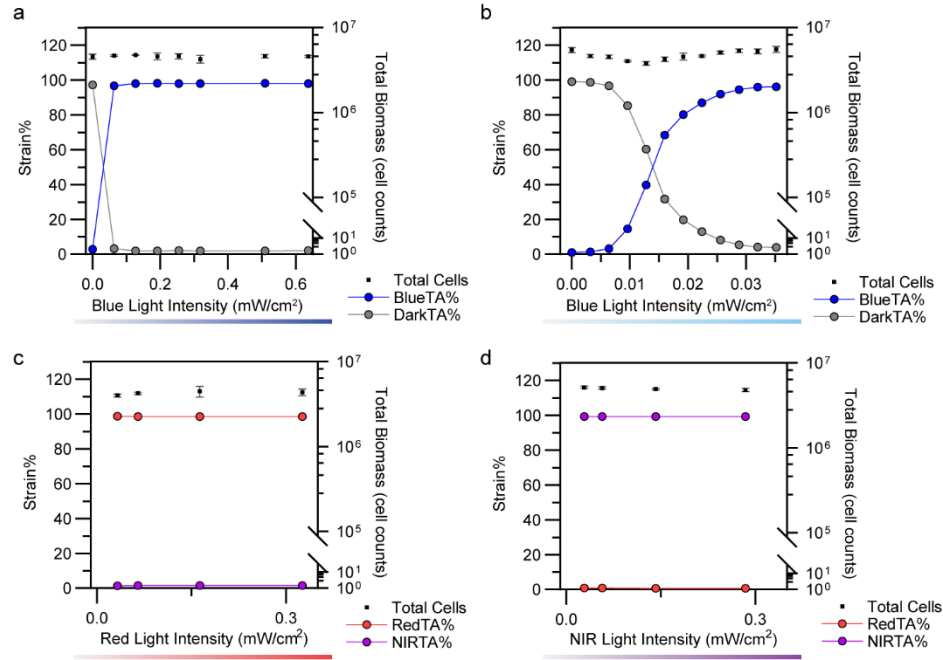

**Supplementary Figure 5.** Effect of modulating intensity on optogeneticTA co-cultures. **a** BlueTA/DarkTA co-culture response to blue light intensity. **b** BlueTA/DarkTA co-culture response to reduced intensity output via placement of the neutral-density (ND9) filters underneath the culture plates. With two ND9 filters, blue light intensities ranging from 0 to 36 $\mu\text{W}/\text{cm}^2$ were accessible, thereby modulating the BlueTA/DarkTA strain population ratio with intensity changes. **c,d** RedTA/NIRTA co-culture response to monochromatic red light (c) or NIR light (d) intensities. All data are shown as mean values with error bars representing standard deviation from $n=3$ biological replicates.

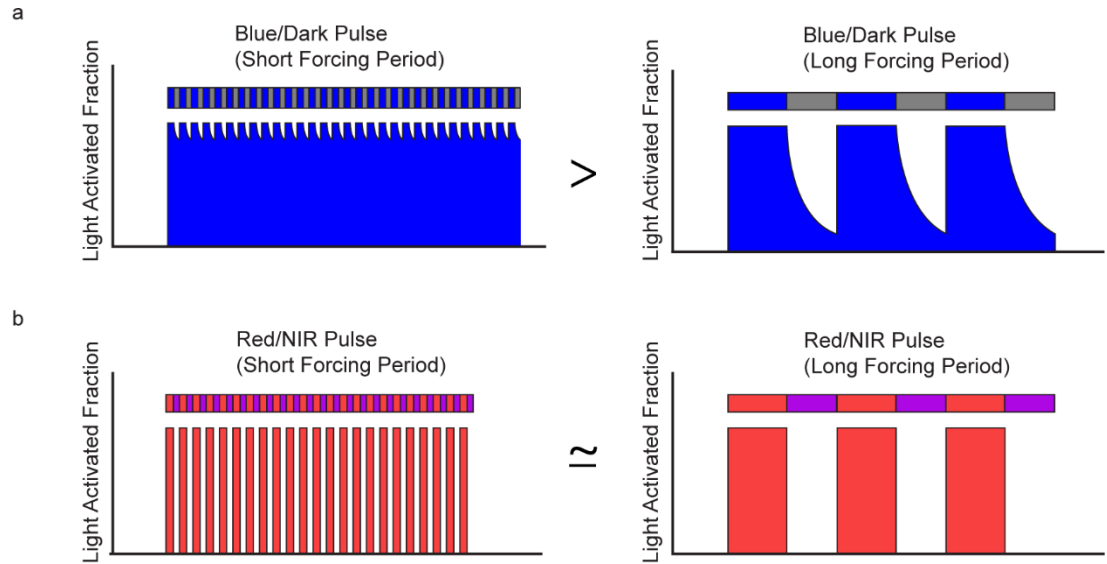

**Supplementary Figure 6.** Simulated light activated fraction when comparing long and short forcing periods at the same dosing percentage. **a** The reversion of YF1-based BlueTA and DarkTA systems following blue light activation is reliant on the dark reversion rate of the photosensitive kinase, YF1 (BlueTA) or YF1<sup>H22P</sup> (DarkTA). Due to the passive deactivation of the blue light-activated systems, the dark periods at short forcing periods ($t_{FP}$) do not persist long enough to fully inactivate blue-light induced kinase/phosphatase activities before the next blue light irradiation, keeping the photoresponsive protein activated throughout<sup>3,17</sup>. **b** In contrast, the red-/NIR-responsive photoresponsive histidine kinases in the RedTA/NIRTA systems can be actively reversed from their red-induced activities using NIR light, and vice versa; thus, it can accelerate the response reversal, rather than depending on slow dark reversion kinetics<sup>13,16</sup>. Red/NIR pulses are hypothesized to have similar activated fractions in long and short forcing periods due to the ability to actively revert red light activation via NIR light.

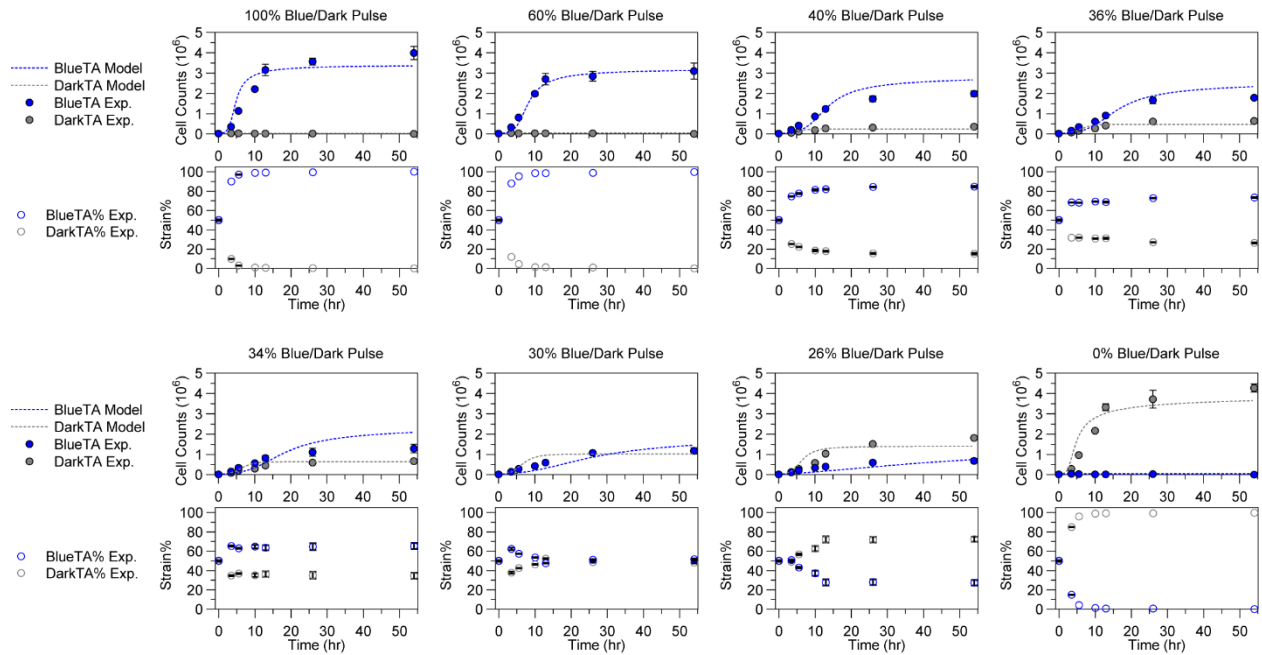

**Supplementary Figure 7.** Trajectories of population BlueTA/DarkTA consortium population distribution under blue light/dark pulsing schedule ($0.32 \text{ mW/cm}^2$, $t_{FP}=2000 \text{ s}$). n% of blue/dark pulse indicates n% of blue light irradiation, followed by (100-n)% of darkness at a certain $t_{FP}$. Top panel shows the trajectory of each strain's cell counts (biomass) at indicated hours while bottom panel shows the strain percentage calculated based on the cell counts. The model fitting the co-culture system is shown as a dotted line. All data are shown as mean values with error bars representing standard deviation from n=3 biological replicates.

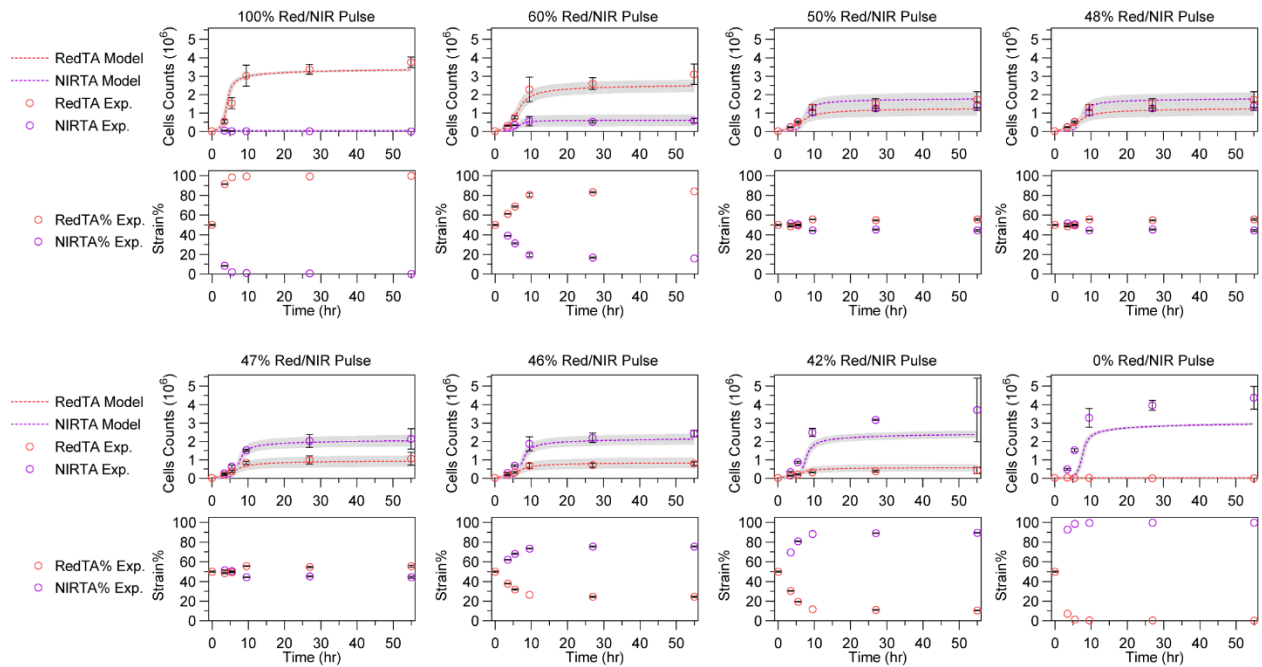

**Supplementary Figure 8.** Trajectories of population composition change of RedTA/NIRTA consortia under red/NIR light pulses ($t_{FP}=2000$ s). n% of Red/NIR pulse indicates n% of red light irradiation, followed by (100-n)% of NIR light irradiation at a certain $t_{FP}$. At fixed red light ($0.17 \text{ mW/cm}^2$) and NIR light ($0.28 \text{ mW/cm}^2$) intensities, alternating red and NIR light irradiation (red/NIR pulsing) was applied to modulate the population composition. Top panels represent the cell counts measured for both RedTA and NIRTA strains while bottom panels show the strain percentage calculated based on cell counts. The model fitting the co-culture system is shown as a dotted line. The grey tubes indicate the confidence intervals of the model trajectories generated based on the initial inoculum biomass uncertainties at 15 uncertainty levels. All data are shown as mean values with error bars representing standard deviation from $n=3$ biological replicates.

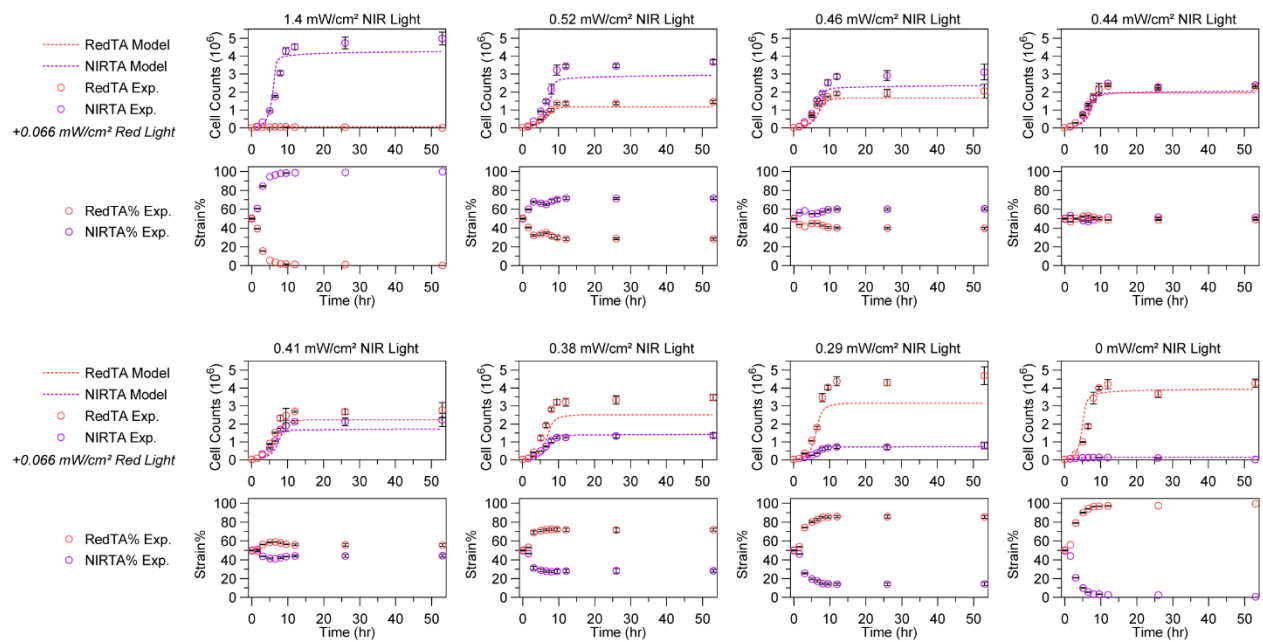

**Supplementary Figure 9.** Trajectories of population composition of RedTA/NIRTA consortia under NIR light intensity changes. Constant red light ($0.066 \text{ mW/cm}^2$) was applied throughout while NIR light intensity was modulated ($0 - 1.4 \text{ mW/cm}^2$) to control the population composition of each strain. Top panels represent the cell counts measured for both RedTA and NIRTA strains while bottom panels show the strain percentage calculated based on the cell counts. The model fitting the co-culture system is shown as a dotted line. All data are shown as mean values with error bars representing standard deviation from $n=3$ biological replicates.

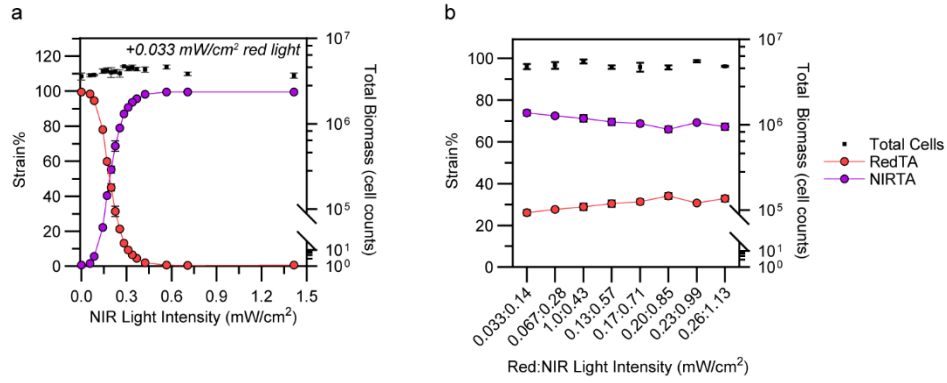

**Supplementary Figure 10.** Effect of light intensity modulation on RedTA/NIRTA consortia. **a** Endpoint population ratio of a RedTA/NIRTA consortium with red light fixed at $0.033 \text{ mW}/\text{cm}^2$ and NIR light intensity screened from 0 – $1.4 \text{ mW}/\text{cm}^2$. **b** Population ratio responses when overall intensities of illumination (red light + NIR light) were varied while keeping the same red:NIR light intensity ratio, keeping equivalent RedTA/NIRTA population ratio throughout the culture. All data are shown as mean values with error bars representing standard deviation from $n=3$ biological replicates.

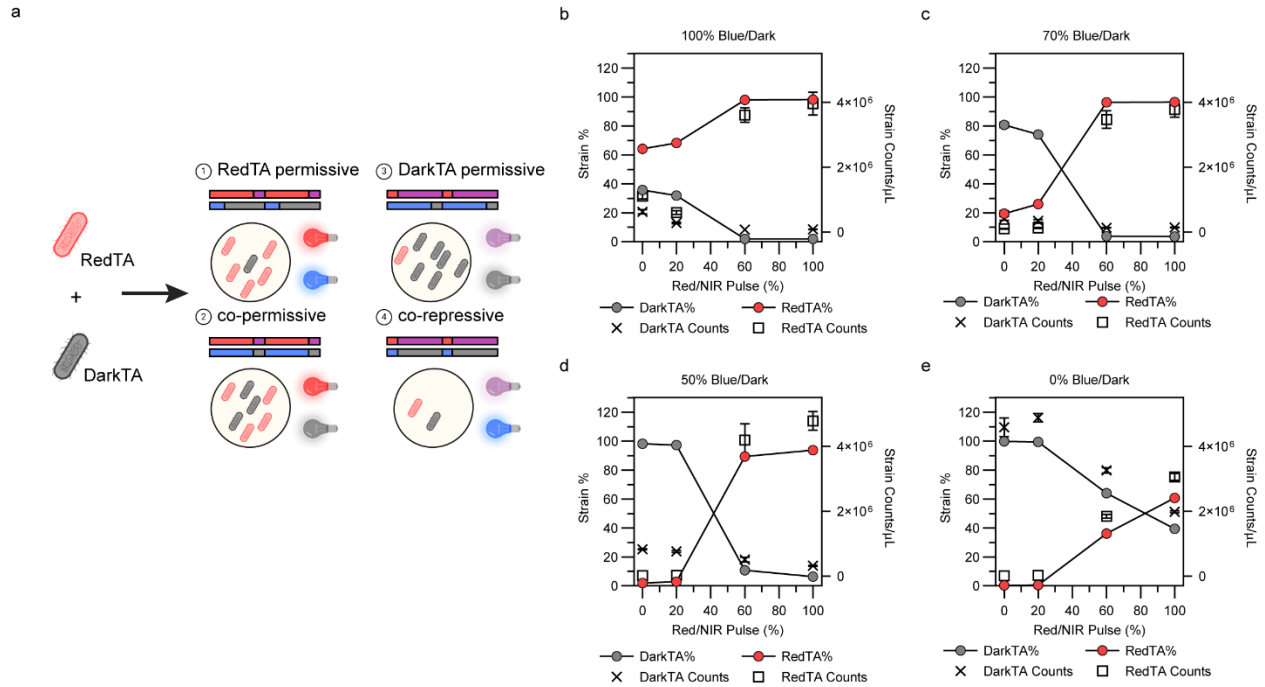

**Supplementary Figure 11.** Two-membered consortia control using RedTA/DarkTA strains. **a** Schematic of RedTA combined with DarkTA at various blue, red, and NIR light schedules and their expected outcome **b-e** Strain percentage and count measurements of RedTA and DarkTA strains in response to various red/NIR pulse ($t_{FP}=2000 \text{ s}$ at $0.17 \text{ mW}/\text{cm}^2$ and $0.28 \text{ mW}/\text{cm}^2$ red and NIR light, respectively) in response to 100% blue/dark (b), 70% blue/dark (c), 50% blue/dark (d), 0% blue/dark (e) at $0.32 \text{ mW}/\text{cm}^2$ blue light. $n\%$ of Red/NIR pulse indicates $n\%$ of red-light irradiation, followed by $(100-n)\%$ of NIR light irradiation at a certain $t_{FP}$, while

blue/dark pulse corresponds to the same scheme with blue light and darkness. Population ratio was measured using flow cytometer after 24 hours of growth at 37 °C, 200 rpm. All data are shown as mean values with error bars representing standard deviation from n=3 biological replicates.

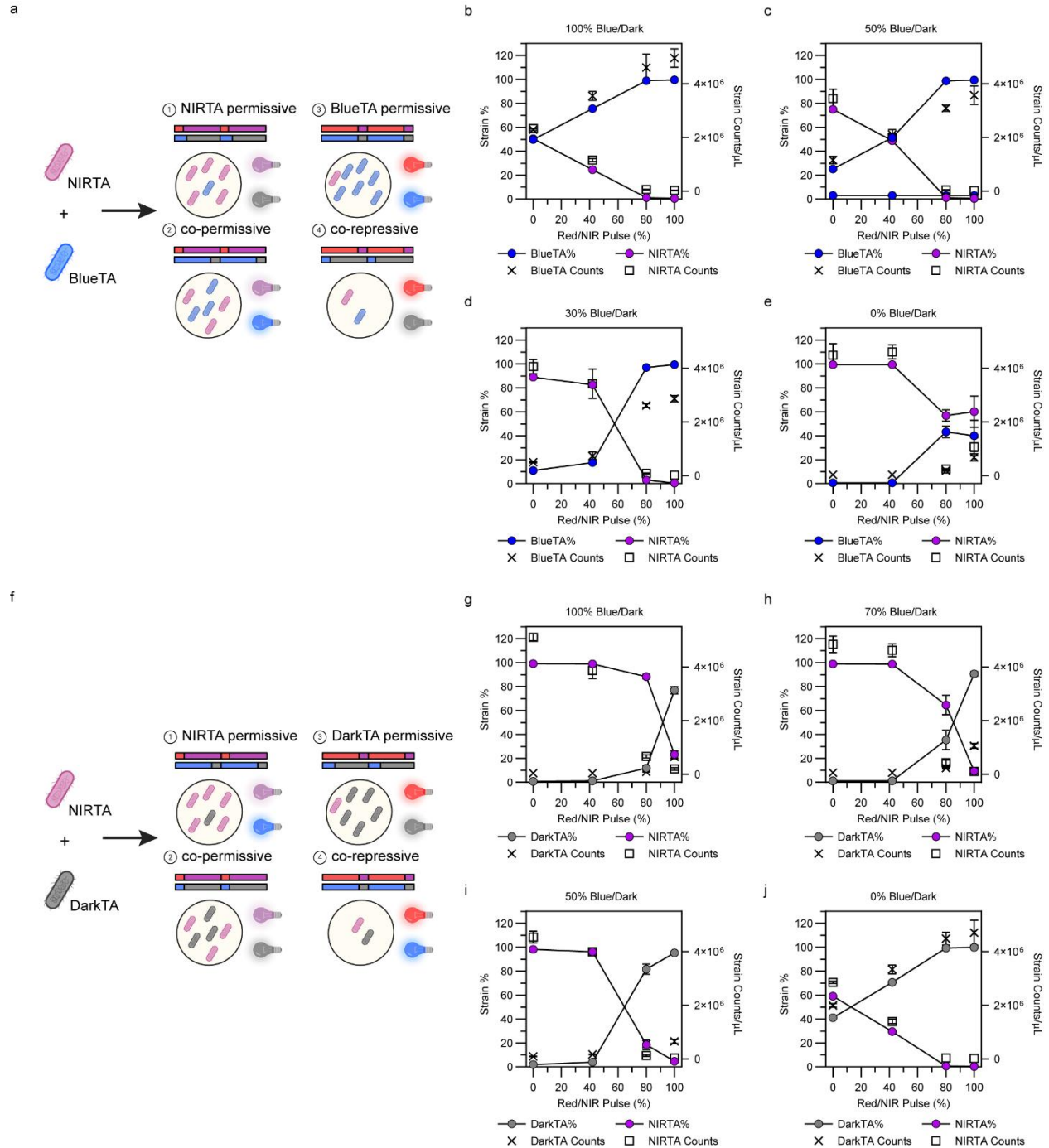

**Supplementary Figure 12.** Two-membered consortia control using NIRTA/BlueTA and NIRTA/DarkTA strains. **a,f** Schematic of NIRTA combined with BlueTA (a) or NIRTA with DarkTA (f) at various blue, red, and NIR light schedules and their expected outcomes. **b-e** Strain

percentage and count measurements of NIRTA and BlueTA strains in response to various red/NIR pulse ($t_{FP}=2000$ s at $0.17 \text{ mW/cm}^2$ and $0.28 \text{ mW/cm}^2$ red and NIR light, respectively) with simultaneous 100% (b), 50% (c), 30% (d), 0% (e) blue/dark pulsing irradiation at $0.32 \text{ mW/cm}^2$ blue light. **g-j** Strain percentage and count measurements of NIRTA and DarkTA strains in response to various red/NIR pulse ($t_{FP}=2000$ s) with simultaneous 100% (g), 70% (h), 50% (i), 0% (j) blue/dark irradiation (same intensities as (b)-(e)). n% of Red/NIR pulse indicates n% of red-light irradiation, followed by (100-n)% of NIR light irradiation at a certain $t_{FP}$, while blue/dark pulse corresponds to the same scheme with blue light and darkness. Population ratio was measured using flow cytometer after 24 hours of growth at $37^\circ\text{C}$, 200 rpm. All data are shown as mean values with error bars representing standard deviation from $n=3$ biological replicates.

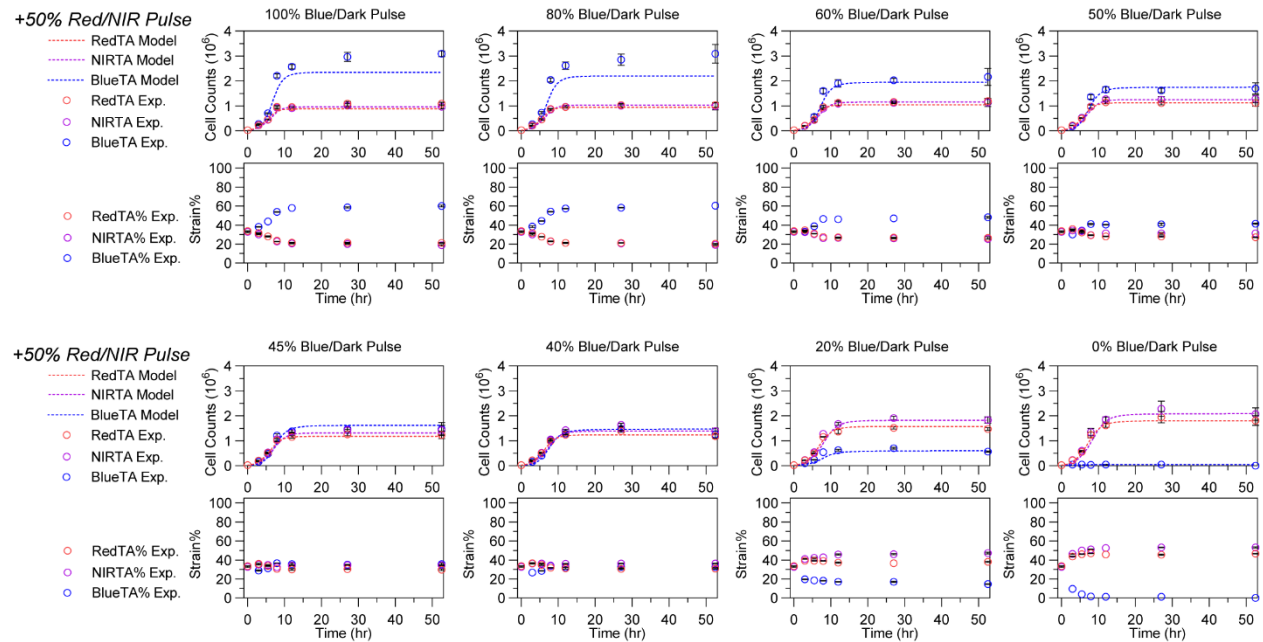

**Supplementary Figure 13.** Three-membered consortia (RedTA, NIRTA, and BlueTA) population trajectories against time. Using fixed red ($0.17 \text{ mW/cm}^2$)/NIR ($0.28 \text{ mW/cm}^2$) pulsing at 50% ($t_{FP}=2000$ s), blue light/dark pulsing was varied (100, 80, 60, 50, 45, 40, 20, and 0%, $t_{FP}=2000$ s). n% of Red/NIR pulse indicates n% of red-light irradiation, followed by (100-n)% of NIR light irradiation at a certain $t_{FP}$, while blue/dark pulse corresponds to the same scheme with blue light and darkness. Top panels represent the cell counts measured for RedTA, NIRTA, and BlueTA strains while bottom panels show the strain percentage calculated based on the cell counts. The mathematical model adopted for three-membered consortia is plotted as a dashed line for each strain. All data are shown as mean values with error bars representing standard deviation from $n=3$ biological replicates.

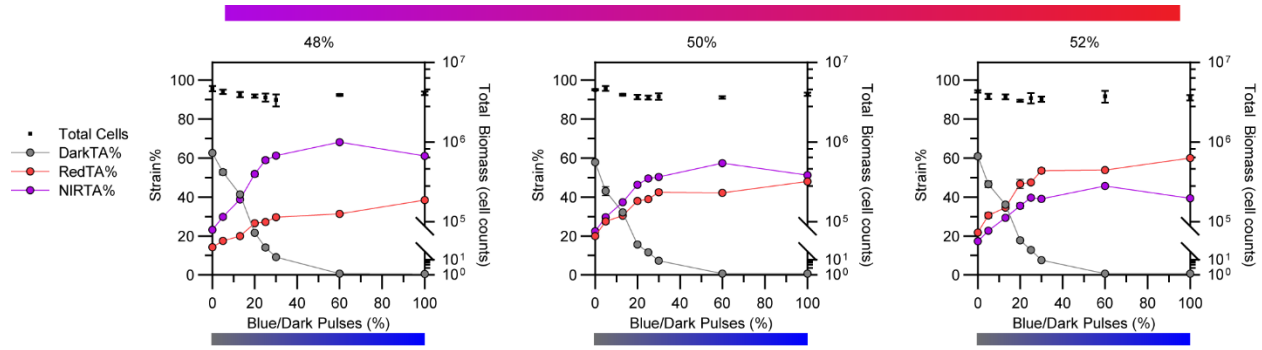

**Supplementary Figure 14.** Three-membered consortia using RedTA, NIRTA, and DarkTA.

Three strains were inoculated at 1:1:1 ratio (final OD: 0.1), followed by irradiation of various blue/dark pulses (x-axis) and red/NIR pulses at 48, 50, and 52% (left to right). n% of Red/NIR pulse indicates n% of red-light irradiation, followed by (100-n)% of NIR light irradiation at a certain $t_{FP}$, while blue/dark pulse corresponds to the same scheme with blue light and darkness. Population ratio was measured using flow cytometer after 48 hours of growth at 37 °C, 200 rpm. In all experiments, blue, red, and NIR light were pulsed at 0.32, 0.17, and 0.28 mW/cm<sup>2</sup>. All data are shown as mean values with error bars representing standard deviation from n=3 biological replicates.

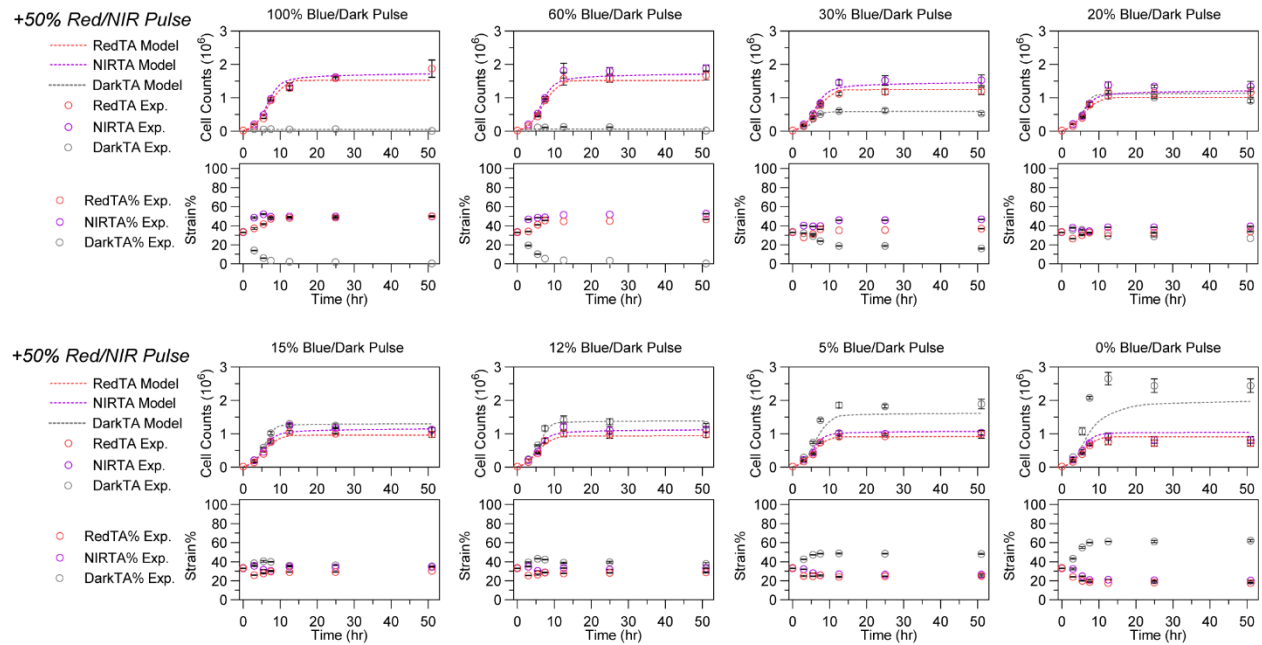

**Supplementary Figure 15.** Three-membered consortia (RedTA, NIRTA, and DarkTA)

population trajectories against time. Using fixed red (0.17 mW/cm<sup>2</sup>)/NIR (0.28 mW/cm<sup>2</sup>) pulsing at 50% ($t_{FP}$=2000 s), blue light (0.32 mW/cm<sup>2</sup>)/dark pulsing was varied (100, 60, 30, 20, 15, 12, 5, and 0%, $t_{FP}$=2000 s). n% of Red/NIR pulse indicates n% of red-light irradiation, followed by (100-n)% of NIR light irradiation at a certain $t_{FP}$, while blue/dark pulse corresponds to the same scheme with blue light and darkness. Top panels represent the cell counts measured for RedTA, NIRTA, and DarkTA strains while bottom panels show the strain percentage calculated based on the cell counts. The mathematical model adopted for three-membered consortia is plotted as a dashed line for each strain. All data are shown as mean values with error bars representing standard deviation from n=3 biological replicates.

a

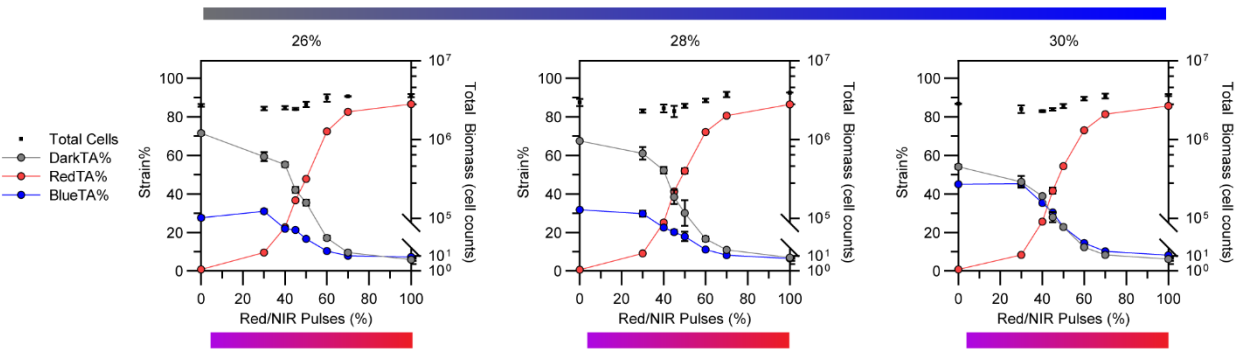

b

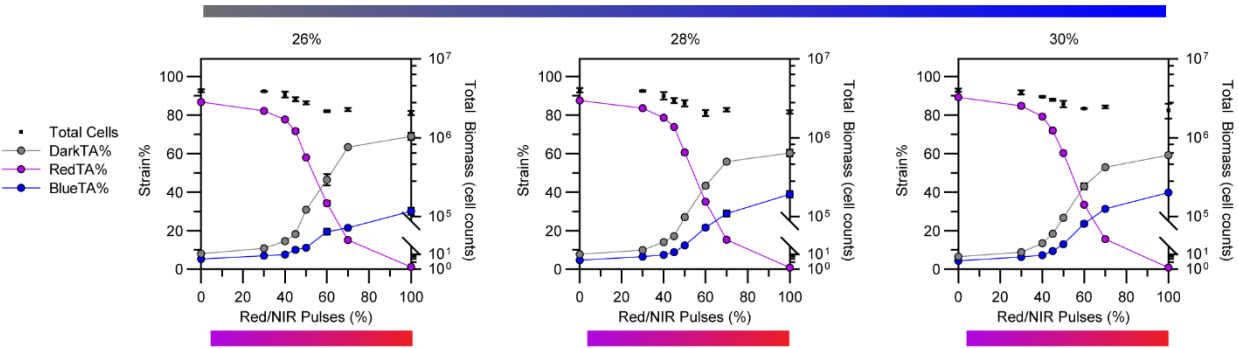

**Supplementary Figure 16.** Three-membered consortia using BlueTA and DarkTA combined with either (a) RedTA or (b) NIRTA. Three strains were inoculated at 1:1:1 ratio (final OD: 0.1), followed by irradiation of various red ($0.17 \text{ mW/cm}^2$)/NIR ($0.28 \text{ mW/cm}^2$) pulses (x-axis) and blue ($0.32 \text{ mW/cm}^2$)/dark pulses at 26, 28, and 30% (left to right). n% of Red/NIR pulse indicates n% of red light irradiation, followed by (100-n)% of NIR light irradiation at a certain $t_{FP}$, while blue/dark pulse corresponds to the same scheme with blue light and darkness. Population ratio was measured using flow cytometer after 48 hours of growth at $37^\circ \text{C}$, 200 rpm. All data are shown as mean values with error bars representing standard deviation from n=3 biological replicates.

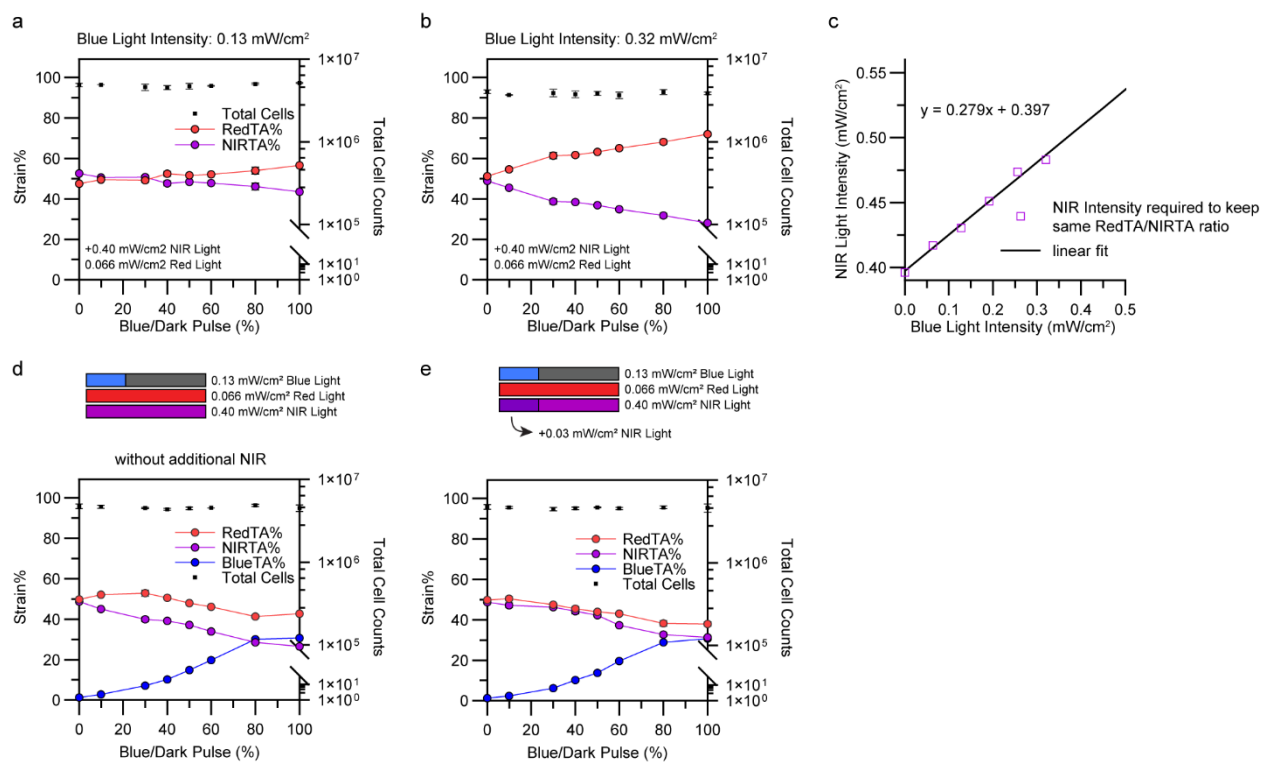

**Supplementary Figure 17.** Effect of blue light on RedTA/NIRTA consortia controlled with intensity modulation. **a,b** RedTA/NIRTA co-culture response to blue/dark pulse (x-axis) using blue light intensity at 0.13 (a) or 0.32 mW/cm<sup>2</sup> (b). NIR light and red light were simultaneously irradiated at 0.40 and 0.066 mW/cm<sup>2</sup> intensities intended to maintain an equivalent ratio of RedTA and NIRTA population. At higher dose of blue light, RedTA composition increased while NIRTA one decreased. **c** Effect of blue light co-irradiation at varying intensities on the RedTA/NIRTA co-culture system was tested by keeping red light intensity constant at 0.066 mW/cm<sup>2</sup> while NIR light intensity was varied to maintain the same RedTA/NIRTA population ratio value (54%:46%, respectively) as in the absence of blue light (Blue Light Intensity = 0). The plot shows a linear relationship between the blue light intensity irradiated versus the NIR intensity required to offset the effect of blue light on the RedTA/NIRTA co-culture. **d,e** The effect of blue light/darkness pulsing on a three-membered consortium comprised of RedTA, NIRTA, and BlueTA. Although blue light pulse (0.13 mW/cm<sup>2</sup>) modulated the BlueTA population, the equivalent ratio of RedTA/NIRTA composition was affected without the NIR offset (d) whereas applying the NIR offset kept the RedTA/NIRTA ratio equivalent. All data are shown as mean values with error bars representing standard deviation from n=3 biological replicates.

a

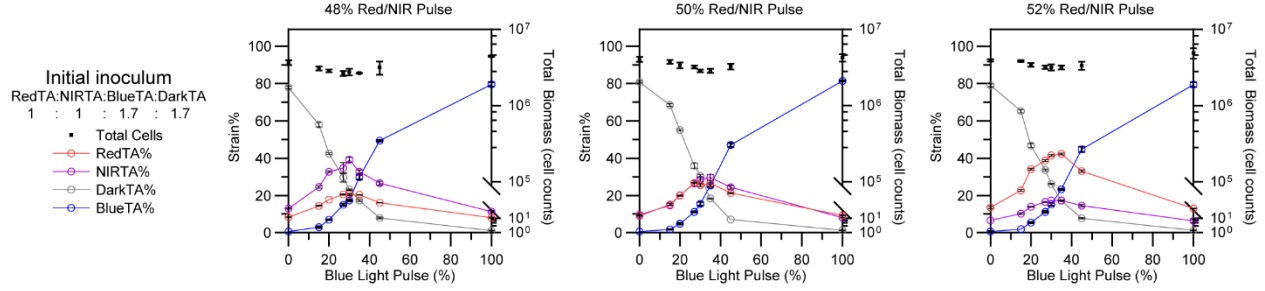

b

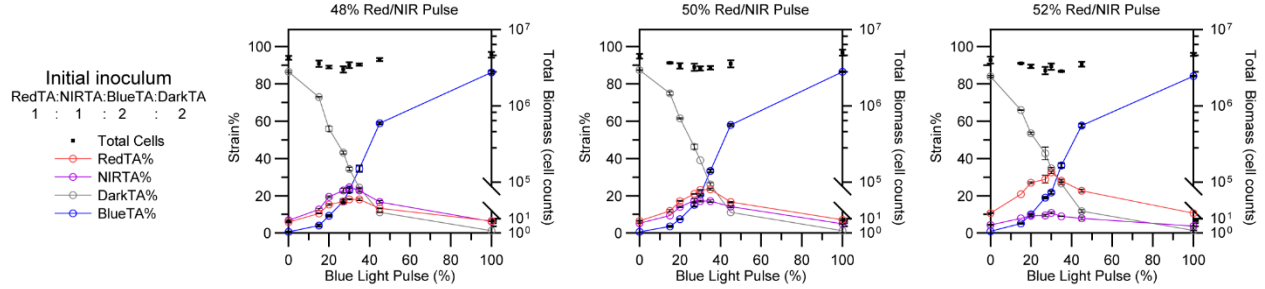

**Supplementary Figure 18.** Optogenetic control over four-membered consortia at various inoculation ratios. **a,b** Endpoint population ratio measurements (post 48 hours) obtained from varying initial inoculum of RedTA:NIRTA:BlueTA:DarkTA at (a) 1:1:1.7:1.7 (b) 1:1:2:2 under irradiation of 48, 50, and 52% red ($0.17 \text{ mW/cm}^2$)/NIR ($0.28 \text{ mW/cm}^2$) pulses (left to right) as a function of varying blue ($0.32 \text{ mW/cm}^2$)/dark pulses (x-axis). n% of Red/NIR pulse indicates n% of red-light irradiation, followed by (100-n)% of NIR light irradiation at a certain $t_{FP}$, while blue/dark pulse corresponds to the same scheme with blue light and darkness. All data are shown as mean values with error bars representing standard deviation from $n=3$ biological replicates.

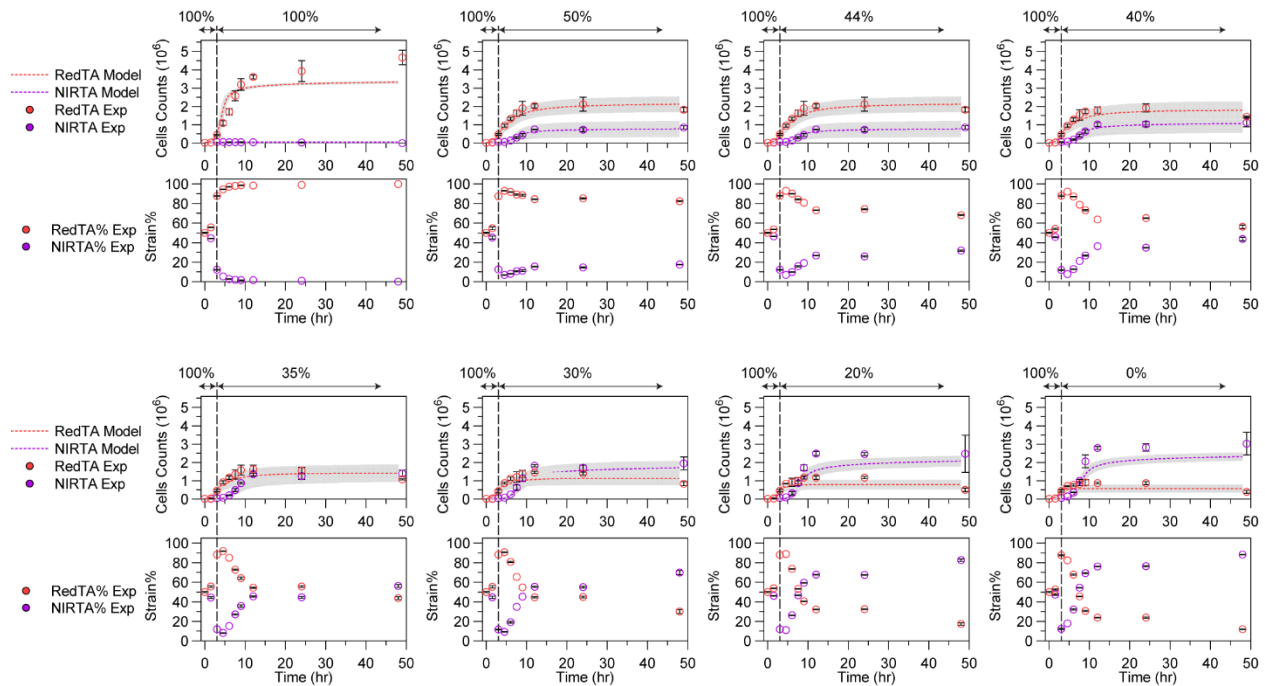

**Supplementary Figure 19.** Dynamic modulation of RedTA/NIRTA consortia composition when varying red/NIR light pulses. Red/NIR pulsing schedule ($t_{FP}=2000$ s) was applied at a fixed red ($0.17 \text{ mW/cm}^2$)/NIR ($0.28 \text{ mW/cm}^2$) intensity to initially populate RedTA for the first three hours (100% pulse), followed by switching red/NIR pulses to various population compositions. Dashed lines indicate the time at which red/NIR pulse was changed. n% of Red/NIR pulse indicates n% of red-light irradiation, followed by (100-n)% of NIR light irradiation at a certain $t_{FP}$. Top panels represent the cell counts measured for RedTA and NIRTA strains while bottom panels show the strain percentage calculated based on the cell counts. The model fitting the co-culture system is shown as a dotted line. The grey tubes indicate the confidence intervals of the model trajectories generated based on the initial inoculum biomass uncertainties at 15 uncertainty levels. All data are shown as mean values with error bars representing standard deviation from $n=3$ biological replicates.

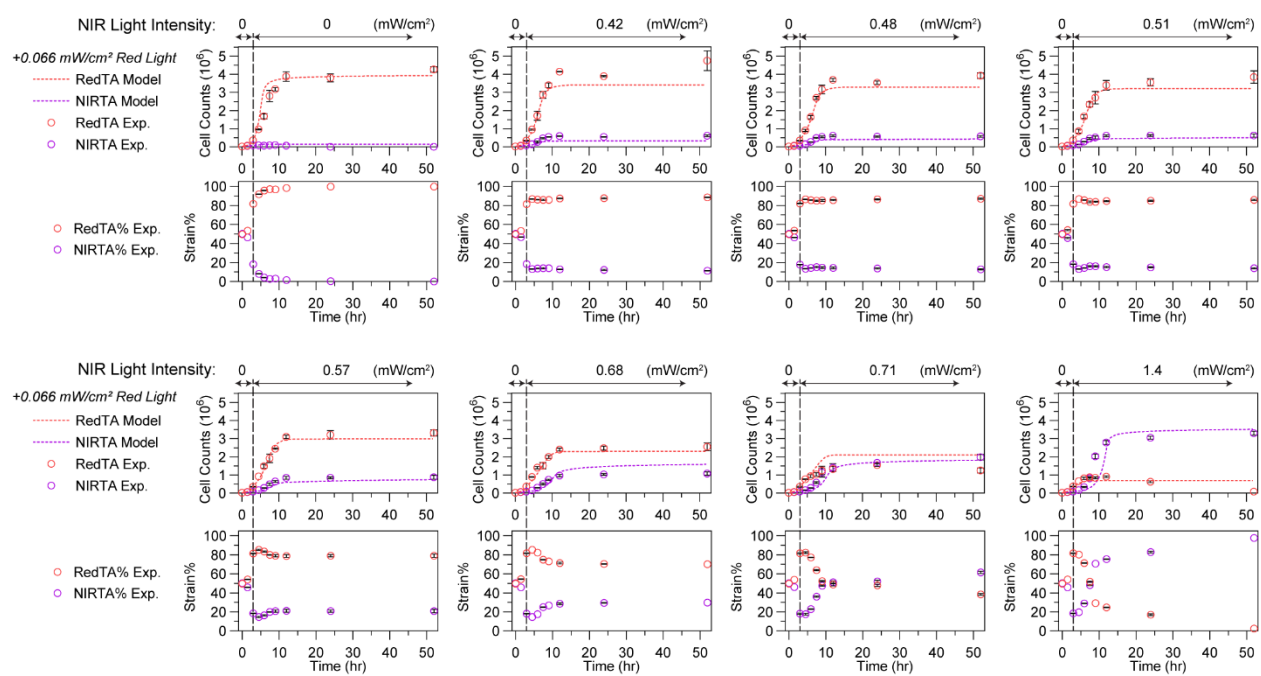

**Supplementary Figure 20.** Dynamic modulation of RedTA/NIRTA consortia composition when varying NIR light intensity. Constant red light ($0.066 \text{ mW/cm}^2$) was applied while NIR light intensity was varied. Light intensity applied for the first three hours enriched RedTA composition (red light: $0.066 \text{ mW/cm}^2$ and NIR Light: $0 \text{ mW/cm}^2$), followed by switching conditions to various NIR light intensities ranging from 0 to $1.4 \text{ mW/cm}^2$ while keeping the red light ($0.066 \text{ mW/cm}^2$) on. Dashed lines indicate the time at which NIR light intensity was changed. Top panels represent the cell counts measured for RedTA and NIRTA strains while bottom panels show the strain percentage calculated based on the cell counts. The model fitting the co-culture system is shown as a dotted line. All data are shown as mean values with error bars representing standard deviation from $n=3$ biological replicates.

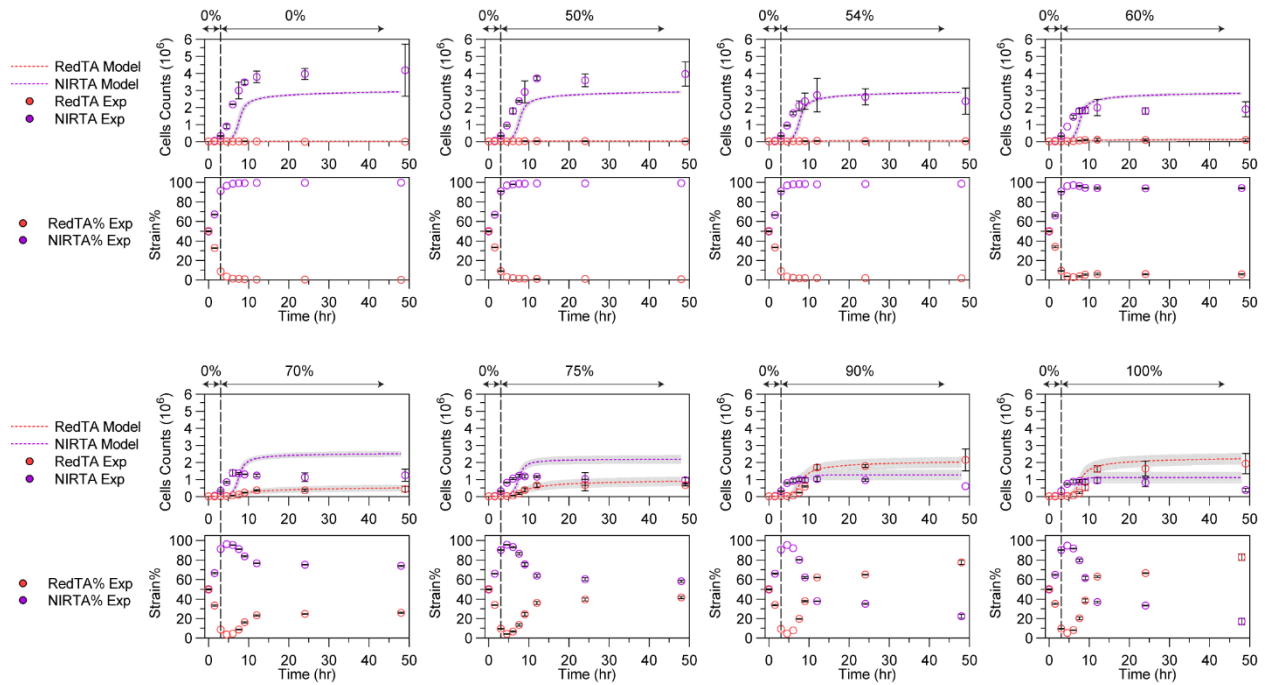

**Supplementary Figure 21.** Dynamic modulation of RedTA/NIRTA consortia composition when varying red/NIR light pulses. At fixed red light ($0.17 \text{ mW/cm}^2$) and NIR light ($0.28 \text{ mW/cm}^2$) intensity applied, red/NIR pulsing schedule ($t_{\text{FP}}=2000 \text{ s}$) was applied to initially populate NIRTA for the first three hours (0% red/NIR pulse, meaning 0 seconds of red light and 2000 seconds of NIR light), followed by switching to various red/NIR pulses. Dashed lines indicate the times at which red/NIR pulses were changed. n% of Red/NIR pulse indicates n% of red-light irradiation, followed by $(100-n)\%$ of NIR light irradiation at a certain $t_{\text{FP}}$. Top panels represent the cell counts measured for RedTA and NIRTA strains while bottom panels show the strain percentage calculated based on the cell counts. The model fitting the co-culture system is shown as a dotted line. The grey tubes indicate the confidence intervals of the model trajectories generated based on the initial inoculum biomass uncertainties at 15 uncertainty levels. All data are shown as mean values with error bars representing standard deviation from $n=3$ biological replicates.

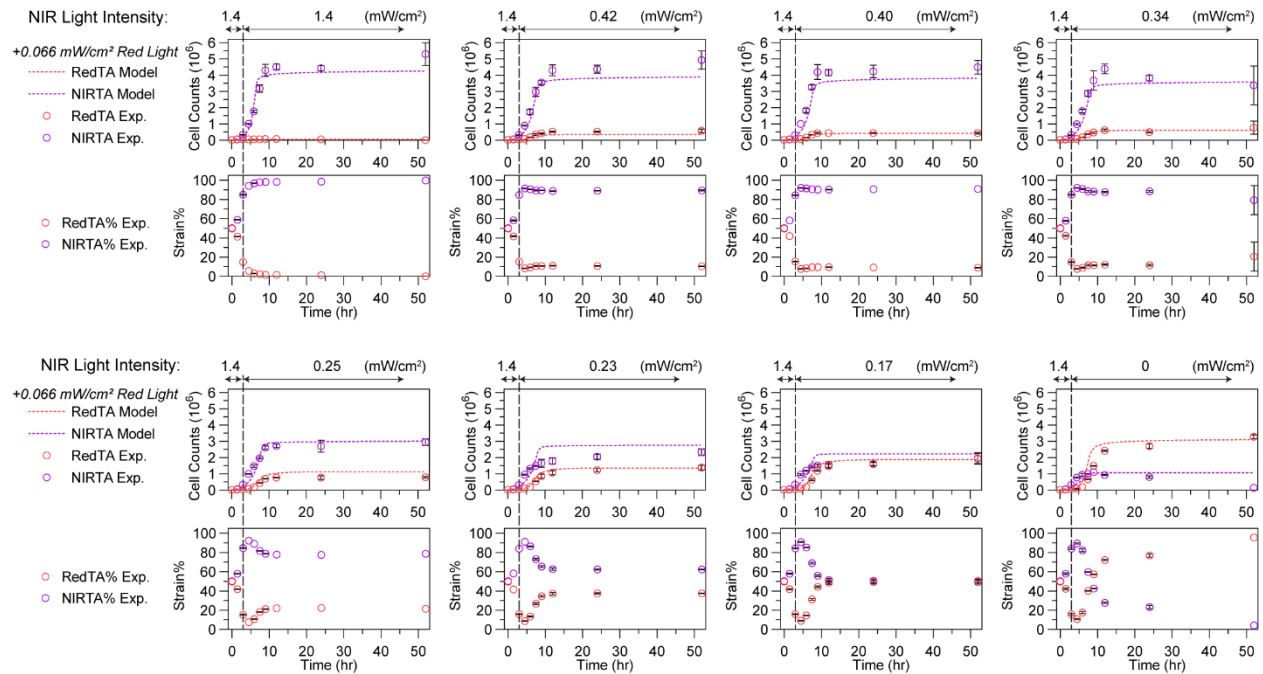

**Supplementary Figure 22.** Dynamic modulation of RedTA/NIRTA consortia composition when varying NIR light intensity. Constant red light (0.066 mW/cm<sup>2</sup>) was applied while NIR light intensity was varied. Light intensity applied for the first three hours enriched NIRTA composition (red light: 0.066 mW/cm<sup>2</sup> and NIR Light: 1.4 mW/cm<sup>2</sup>), followed by switching to various NIR intensities at 3 hours (vertical dashed line). Top panels represent the cell counts measured for RedTA and NIRTA strains while bottom panels show the strain percentage calculated based on the cell counts. The model fitting the co-culture system is shown as a dotted line. All data are shown as mean values with error bars representing standard deviation from n=3 biological replicates.

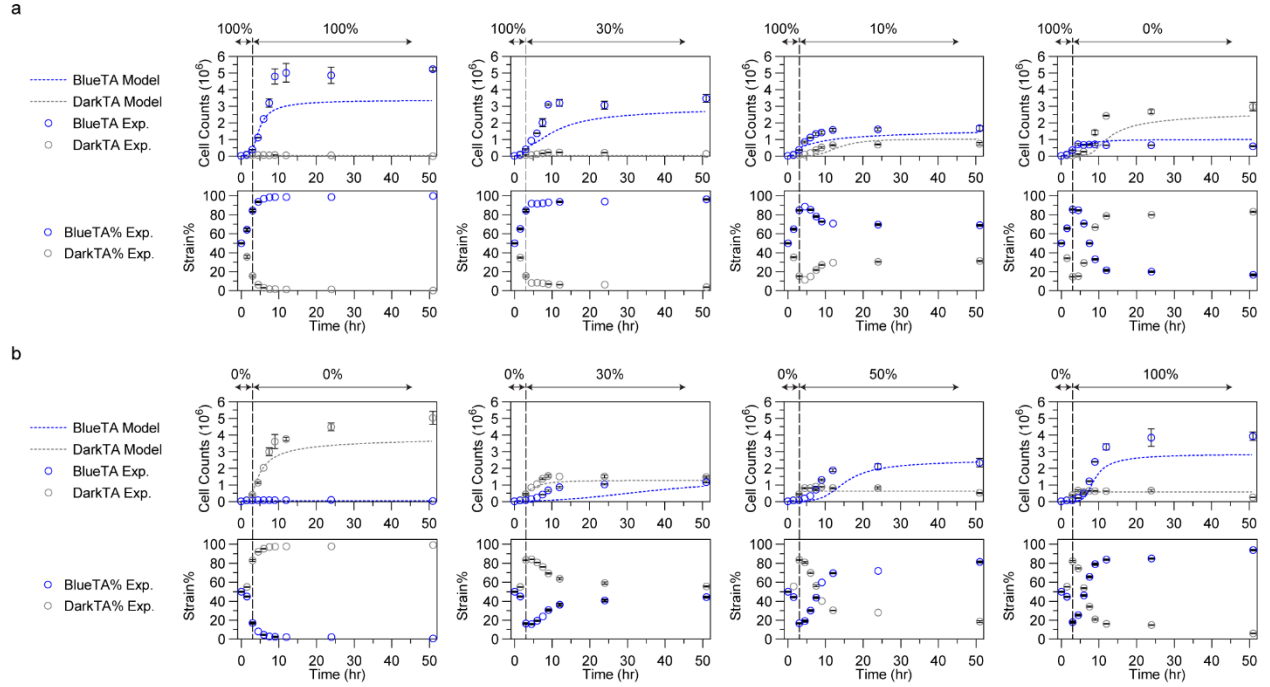

**Supplementary Figure 23.** Dynamic modulation of BlueTA/DarkTA consortia composition using blue light/dark pulses ($0.32 \text{ mW/cm}^2$, $t_{FP}=2000 \text{ s}$). n% of blue/dark pulse indicate n% of blue light irradiation, followed by (100-n)% of darkness at a certain $t_{FP}$. Top panels show the cell counts measured of each strain's trajectories over time while the bottom panels show the strain percentages calculated from the cell counts. Vertical dashed lines indicate the 3-hour point where the light conditions were changed. Mathematical model adopted for the BlueTA/DarkTA co-culture was fit (dotted lines) to the experimental data (circle). All data are shown as mean values with error bars representing standard deviation from n=3 biological replicates.

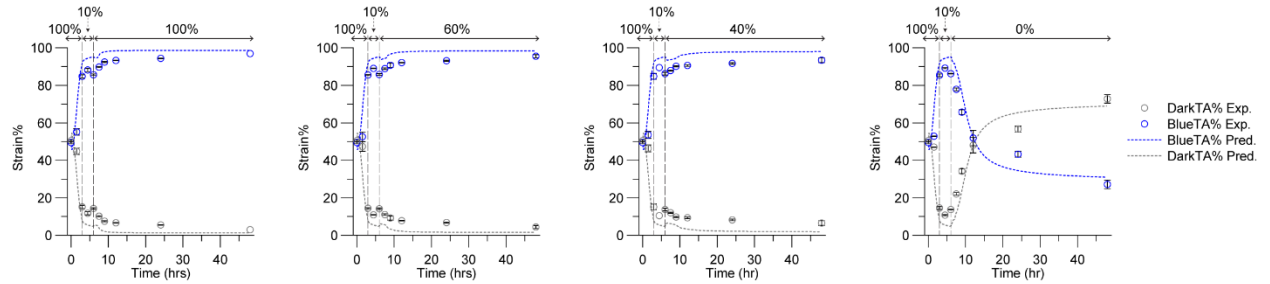

**Supplementary Figure 24.** Model validation of BlueTA/DarkTA co-culture dynamics under novel light schedules. Co-culture was exposed to 100% blue light/dark pulse (at $0.32 \text{ mW/cm}^2$ blue light) for 3 hours, then 10% blue/dark pulse for the next 3 hours, followed by the indicated pulsing conditions. n% of blue/dark pulse indicates n% of blue light irradiation, followed by (100-n)% of darkness at a certain $t_{FP}$. Model predicted trajectories are indicated in dotted lines and experimental validation data points are in circles. Vertical dashed lines indicate the times at which the light pulses were changed (3 and 6 hours). All data are shown as mean values with error bars representing standard deviation from n=3 biological replicates.

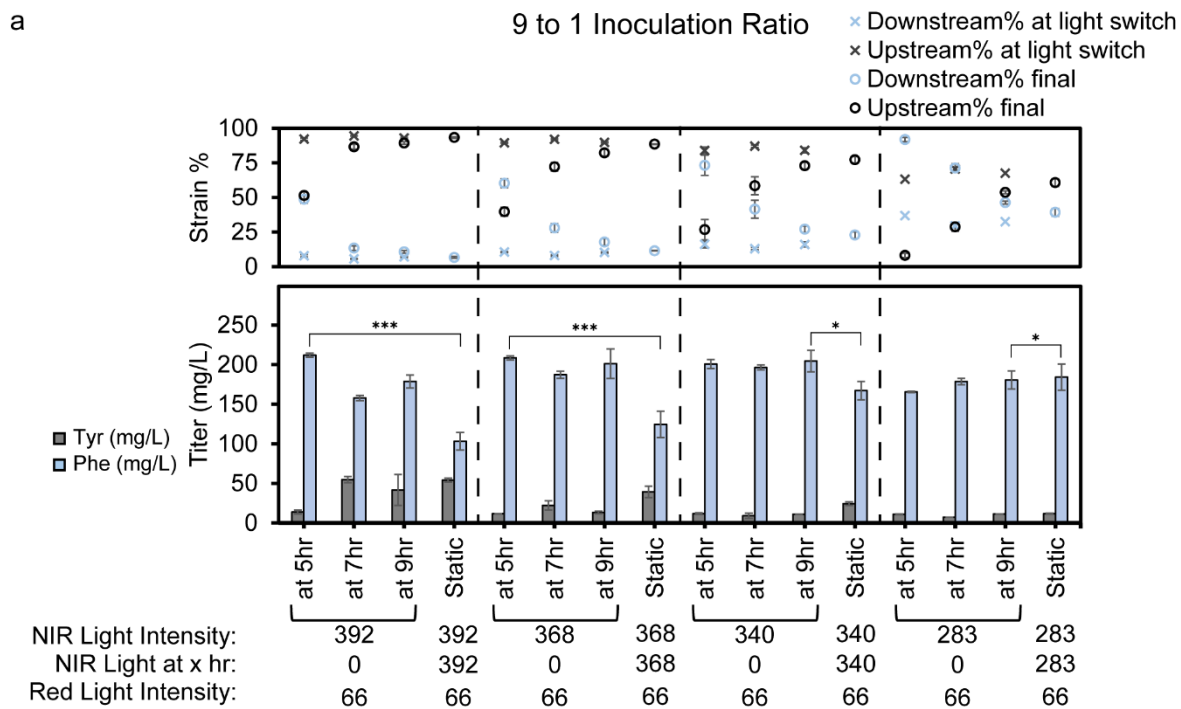

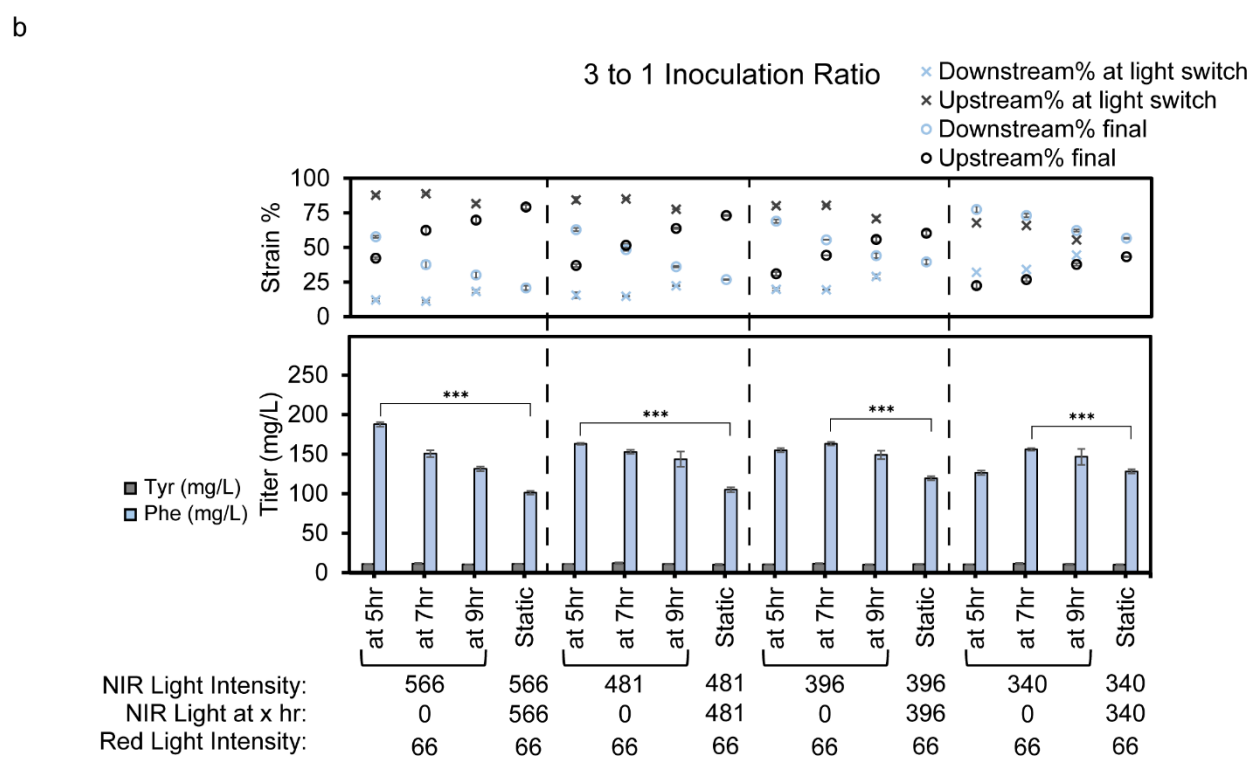

**Supplementary Figure 25.** Dynamic control of phenol production using the optogeneticTA system by varying inoculation ratio between 9-to-1 (a) and 3-to-1 (b) while modulating light

switching times. Tyrosine-producing upstream strain was controlled using the NIRTA system while the phenol-producing downstream strain was controlled using the RedTA system. Co-cultures were irradiated with red light at 66 $\mu\text{W}/\text{cm}^2$ and indicated NIR light intensities for 5, 7, or 9 hours before switching the NIR light to 0 $\mu\text{W}/\text{cm}^2$ while keeping the red light on at 66 $\mu\text{W}/\text{cm}^2$ for the rest of the cultivation. Static indicates no change in NIR light intensity until the end of cultivation. All phenol and tyrosine titers are indicated in means (mg/L) and the error bars represent standard deviation. Strain percentages are calculated as mean from strain percentages obtained from cell counts on the flow cytometer and the error bars represent standard deviation. Statistics are derived from t-test (\*\*\*$P < 0.001$, \*\*$P < 0.002$, \*$P < 0.033$, ns: not significant).

**Supplementary Figure 26.** Dynamic control of phenol production using the optogeneticTA system at 1-to-1 inoculation ratio while modulating light switching times. See Supplementary Fig. 25 for description. All phenol and tyrosine titers are indicated in means (mg/L) and the error bars represent standard deviation. Strain percentages are calculated as mean from strain percentages obtained from cell counts on the flow cytometer and the error bars represent standard deviation. Statistics are derived from t-test (\*\*\*$P < 0.001$, \*\*$P < 0.002$, \*$P < 0.033$, ns: not significant).
